# Weak Acid Resistance A (WarA), a novel transcription factor required for regulation of weak-acid resistance and spore-spore heterogeneity in *Aspergillus niger*

**DOI:** 10.1101/788141

**Authors:** Ivey A. Geoghegan, Malcolm Stratford, Mike Bromley, David B. Archer, Simon V. Avery

## Abstract

Propionic, sorbic and benzoic acids are organic weak acids that are widely used as food preservatives, where they play a critical role in preventing microbial growth. In this study, we uncovered new mechanisms of weak acid resistance in moulds. By screening a library of 401 transcription-factor deletion strains in *Aspergillus fumigatus* for sorbic acid hypersensitivity, a previously uncharacterised transcription factor was identified, and named as WarA (Weak Acid Resistance A). The orthologous gene in the spoilage mould *Aspergillus niger* was identified and deleted. WarA was required for resistance to a range of weak acids, including sorbic, propionic and benzoic acids. A transcriptomic analysis was performed to characterise genes regulated by WarA during sorbic acid treatment in *A. niger*. Several genes were significantly upregulated in the wild type compared with a Δ*warA* mutant, including genes encoding putative weak acid detoxification enzymes and transporter proteins. Among these was An14g03570, a putative ABC-type transporter which we found to be required for weak acid resistance in *A. niger*. We also show that An14g03570 is a functional homologue of the *Saccharomyces cerevisiae* protein Pdr12p, and therefore named as PdrA. Lastly, resistance to sorbic acid was found to be highly heterogeneous within genetically-uniform populations of ungerminated *A. niger* conidia, and we demonstrate that *pdrA* is a determinant of this heteroresistance. This study has identified novel mechanisms of weak acid resistance in *A. niger* which could help to inform and improve future food spoilage prevention strategies.

**IMPORTANCE:** Weak acids are widely used as food preservatives, as they are very effective at preventing growth of most species of bacteria and fungi. However, some species of moulds can survive and grow in the concentrations of weak acid employed in food and drink products, thereby causing spoilage with resultant risks for food security and health. Current knowledge of weak acid resistance mechanisms in these fungi is limited, especially in comparison to that in yeasts. We characterised gene functions in the spoilage mould species *Aspergillus niger* which are important for survival and growth in the presence of weak acid preservatives. Such identification of weak acid resistance mechanisms in spoilage moulds will help to design new strategies to reduce food spoilage in the future.

## INTRODUCTION

Microbiological spoilage of food and drinks is a serious threat to food security and human health. It is estimated that 25% of global food produced annually is lost due to contamination and degradation by microorganisms (1). Such spoilage also directly imperils human health, for example due to the production of toxins by the microorganism. Preventing microbial spoilage is therefore a key to safeguarding food supply and safety. The use of chemical food preservatives to inhibit growth of bacteria and fungi is a ubiquitous and generally effective strategy in reducing spoilage (2). Some of the most commonly used preservatives are weak organic acids such as propionic, sorbic and benzoic acids. These are usually included in food and drink products in the form of calcium, potassium and sodium salts, respectively.

Weak acid preservatives are broad-spectrum antimicrobials that directly inhibit the growth of yeasts, moulds and bacteria. Although their precise mechanism of action has not yet been fully determined, it is known that weak acid preservatives cause a reduction of cytoplasmic pH (3, 4), and inhibit nutrient uptake (5, 6). It is also known that weak acids tend to be fungistatic rather than fungicidal, especially at the levels legally permitted in food and drinks. In most cases, microbial growth is completely inhibited by the weak acid levels used in food and drink products. However, certain species of yeasts and moulds demonstrate elevated resistance to weak acids, and are therefore capable of causing food spoilage (7, 8).

Weak acid resistance can be attributed in part to the enzymatic degradation of certain acids (e.g., benzoic, sorbic and cinnamic acids), which nullifies their antimicrobial effects. Benzoate can be catabolized through a pathway involving an initial hydroxylation step. The enzyme responsible (benzoate *para*-hydroxylase) has been found to be required for resistance to benzoic acid in *Aspergillus niger* and *A. nidulans* (9, 10). Sorbic and cinnamic acids (and certain other structurally related acids) are degraded by decarboxylation (11). In moulds such as *A. niger*, a cluster of three genes is required for this process, encoding a transcription factor (SdrA), a decarboxylase (CdcA; formerly OhbA1 or FdcA) and a prenyltransferase (PadA) (12–15). Deletion of any of these genes reduces, but does not eliminate, resistance to sorbic acid. Thus, additional and as-yet uncharacterized mechanisms of weak acid resistance must operate in this mould species. Enzymatic decarboxylation of weak acids also occurs in numerous yeast species (16). However, contrary to the case in moulds, deletion of the phenylacrylic acid decarboxylase gene (*PAD1*) in the yeast *S. cerevisiae* does not decrease weak acid resistance (16). Furthermore, certain spoilage yeasts do not appear to decarboxylate weak acids at all, suggesting that alternative mechanisms of resistance also operate in these species.

Mechanisms of weak acid resistance have been best characterized in *S. cerevisiae*. One of the key genes required for resistance is *PDR12*, encoding an ATP-Binding Cassette (ABC) transporter (17). *PDR12* is required for resistance to carboxylic acids with chain lengths between 1 and 7, proposedly by mediating the efflux of weak acid anions from the cell in an energy-dependent manner (18). *PDR12* is itself transcriptionally regulated by War1p, a Zn2Cys6 zinc finger transcription factor that binds to weak acid response elements (WARE) in the *PDR12* promoter (19). Another transcription factor, Haa1p, is also required for resistance to weak acids in *S. cerevisiae*, by regulating the transcription of membrane multidrug transporters (Tpo2p and Tpo3p) amongst other, less well characterized genes (20). High-throughput mutant screens have helped to identify many other genes which influence weak acid resistance in *S. cerevisiae*. For example, Mollapour et al. (21) reported 237 genes which were required for wild-type resistance to sorbic acid, and a further 34 which resulted in enhanced sorbic acid resistance when deleted. A similar study, also in *S. cerevisiae,* revealed 650 determinants of acetic acid resistance (22). Unfortunately, there is a distinct lack of equivalent data in any other fungal species, including moulds. Considering the propensity of mould fungi to cause food spoilage, understanding the genetic determinants of weak acid resistance in these species is very important.

An additional and historically-overlooked determinant of antimicrobial resistance is the phenotypic heterogeneity that exists within microbial cell populations. Phenotypic heterogeneity is a phenomenon observed within isogenic cell populations, whereby individual cells can display a markedly different phenotype despite being genetically identical. This has been recognised as an important determinant of microbial cell survival in response to antimicrobial agents and other environmental stressors (23–25). Phenotypic heterogeneity in weak acid resistance (heteroresistance) has been found in cell populations of *S. cerevisiae* and the spoilage yeast *Zygosaccharomyces bailli* (8, 26, 27). However, there has been no investigation to date of whether weak acid resistant subpopulations exist in populations of mould spores, although heterogeneity is known to arise in *A. niger* spore populations as a consequence of asynchronous conidial maturation (28). The presence of weak acid resistant spore subpopulations could have significant implications for spoilage control strategies and is therefore worthy of investigation.

In this study, we report the identification and characterization of a novel transcription factor (Weak Acid Resistance A), that is required for resistance to weak acid preservatives in *A. niger* and *A. fumigatus*. Furthermore, we identify and characterize genes that are putatively regulated by WarA, including a gene encoding a putative membrane transporter protein with similarity to *S. cerevisiae* Pdr12p and which, we show, mediates weak acid resistance and heteroresistance in *A. niger*. These data significantly enhance our understanding of weak acid resistance in moulds, and highlight both similarities and differences in weak acid resistance strategies between yeast and mould fungi. (This article was submitted to an online preprint archive (29))

## RESULTS

### A screen for transcription-factor deletion strains sensitive to sorbic acid identifies *warA*

To find genes associated with weak acid resistance, an *Aspergillus fumigatus* transcription-factor deletant collection (30) was screened for sorbic acid sensitivity. *Aspergillus fumigatus* is not commonly associated with food spoilage, and displays relatively high sensitivity to weak acids such as sorbic acid (14). However, deletant collections are available in *A. fumigatus*, unlike Aspergilli associated with food spoilage. It was reasoned that transcription factors associated with weak acid resistance in *A. fumigatus* may be conserved in related spoilage species such as *A. niger*. This resource comprised a library of 401 deletion strains of non-essential transcription factors. To determine sensitivity of the deletion strains, radial growth was compared on agar medium with and without sorbic acid (Fig. 1). This revealed two deletion strains which were highly sensitive to sorbic acid compared with the wild-type strain (Δ*metR*, and Δ*AFUB_000960*) (Fig. 1 and Fig. 2A). A number of other strains exhibited moderate sensitivity to sorbic acid, including Δ*creA* (Δ*AFUB_027530*), Δ*devA* (Δ*AFUB_030440*) (Fig. 1B and Fig. 2A), Δ*rfeD* (*ΔAFUB_022280*), Δ*AFUB_020350* and Δ*AFUB*_*054360* (Fig. 1B).

**FIG 1.**
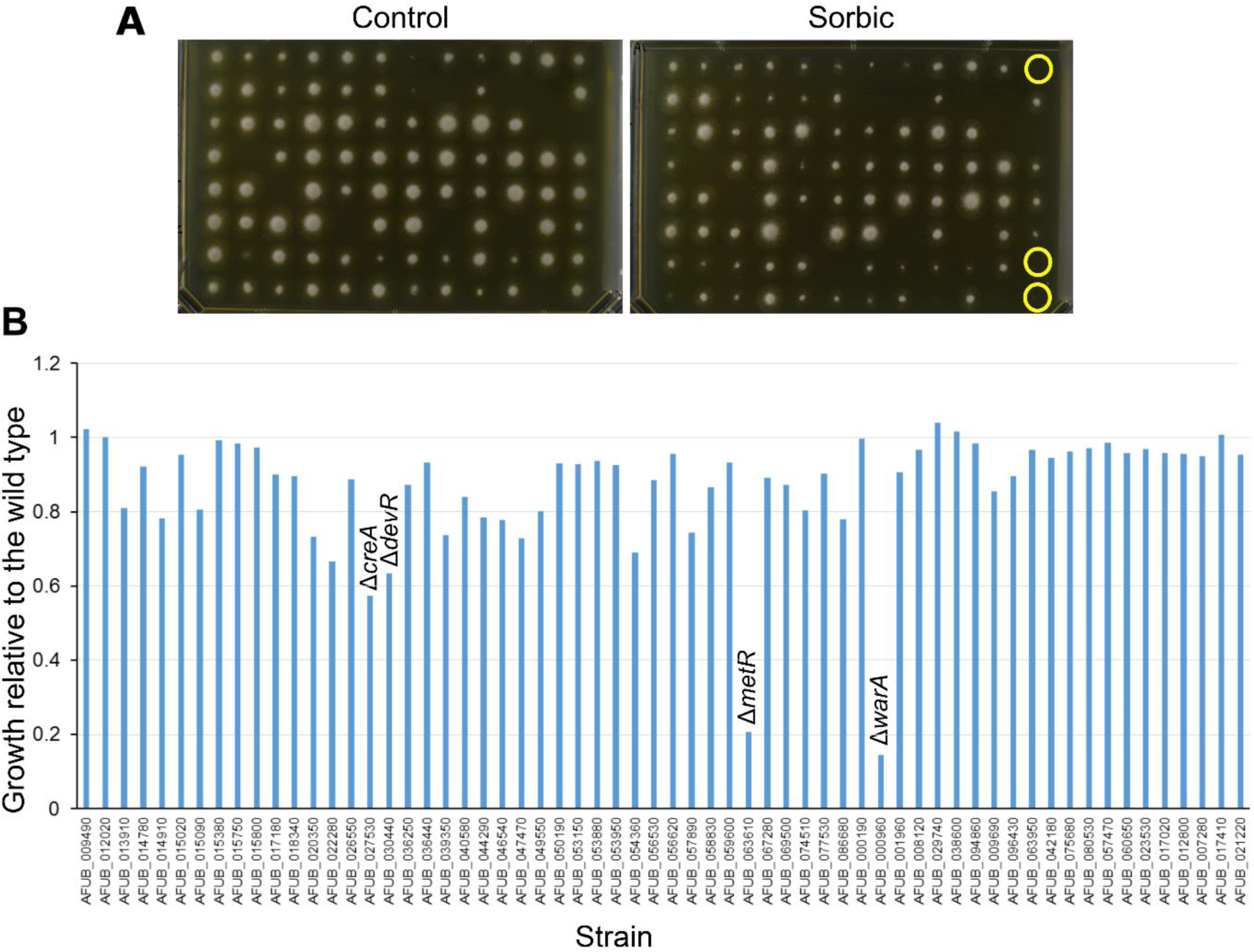
Screening of *A. fumigatus* deletion library. (A) Example of *A. fumigatus* deletant library screen. Conidial suspensions of the different deletants were arrayed in 96-well plates, and transferred to growth medium using a 96-pin tool. Examples of putatively sorbic acid sensitive strains are circled in yellow. (B) Sensitivity of *A. fumigatus* transcription factor deletion strains to sorbic acid. 62 strains were identified from the initial screen in (A) as putatively sorbic acid hypersensitive, and subjected to a second round of screening as outlined in Materials and Methods. Sensitivity to sorbic acid relative to the WT strain is shown (a value of 1 indicates identical sensitivity of the deletion strain to the WT, according to radial growth). “Δ*warA”* refers to Δ*AFUB_000960*.

**FIG 2.**
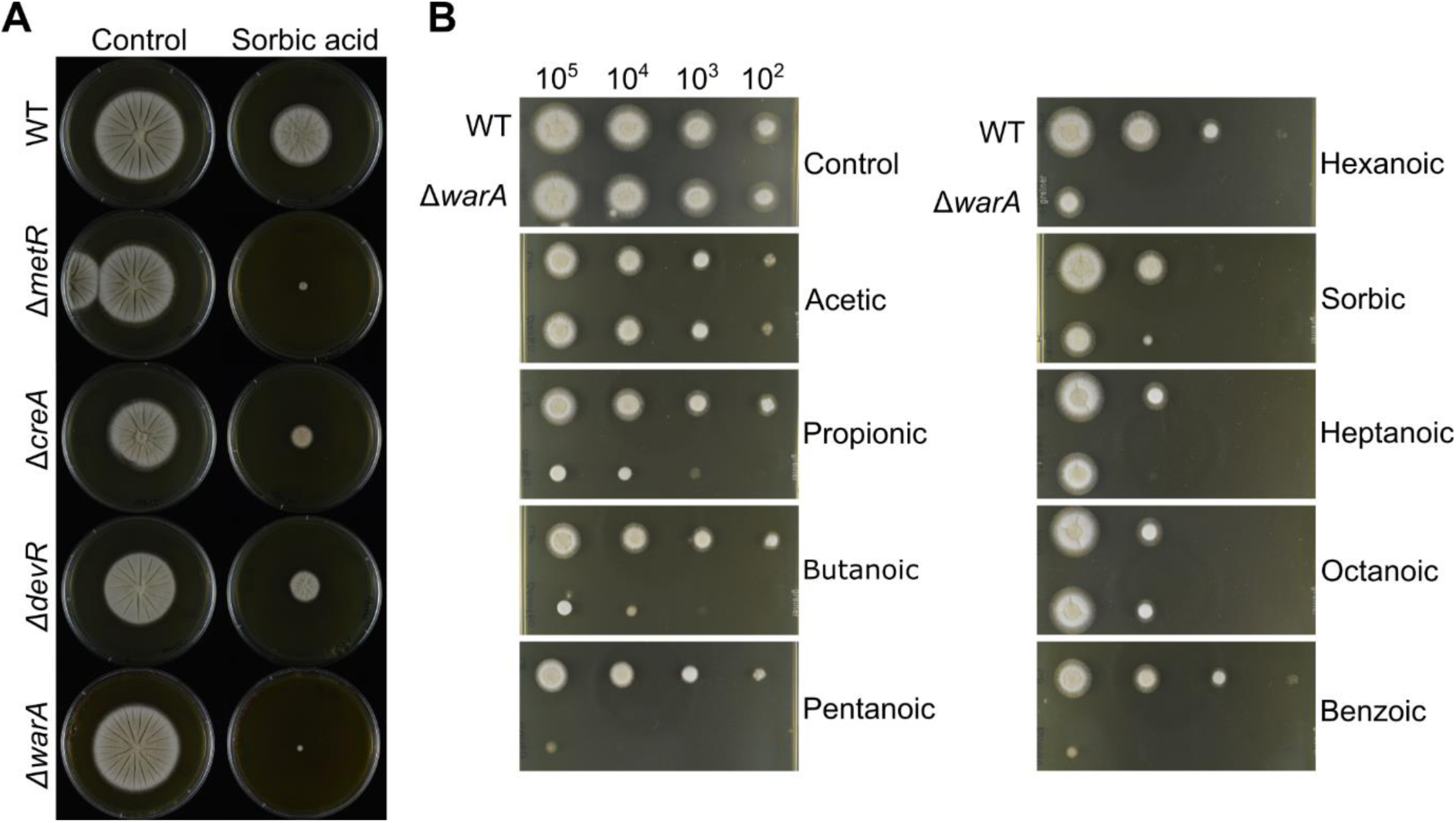
Growth of *A. fumigatus* transcription factor deletion strains on medium containing weak acids. (A) Radial growth of *A. fumigatus* transcription factor deletion strains on agar containing 0.5 mM sorbic acid. Images were captured after 3 d growth at 37°C. (B) Radial growth of *A. fumigatus* Δ*warA* and wild-type on agar containing weak acids. Plates were inoculated with a 10-fold dilution series of conidial suspensions; approximate numbers of conidia are indicated above the pictures. Images were captured after 2 d growth at 28°C, and are representative of 2-3 independent experiments. Concentrations of acids used are given in the Materials and Methods.

MetR is a bZIP-type transcription factor mediating transcriptional regulation of genes involved in sulphur uptake and utilization (31). Because sorbic acid is known to decrease cellular uptake of some nutrients (6, 32), it was hypothesised that sulphur limitation could be a cause of sorbic acid sensitivity in the Δ*metR* deletion strain. To test this, sorbic acid sensitivity of the Δ*metR* strain was determined in medium supplemented with the sulphur-containing amino acid methionine. This showed that the sorbic acid sensitivity of the Δ*metR* strain was abolished in the presence of supplementary methionine (Fig. S1).

*AFUB_000960* encodes a Zn2Cys6-type transcription factor which contains a fungal specific transcription factor domain (PF11951). To further investigate the role of *AFUB_000960* in weak acid resistance, sensitivity of the deletion strain to a range of weak acids was evaluated (Fig. 2B). The Δ*AFUB_000960* strain was sensitive to propionic, butanoic, pentanoic, hexanoic, sorbic and benzoic acids, but not to acetic acid. Because of these phenotypes, *AFUB_000960* was named *Weak Acid Resistance A* (*warA)*.

### A WarA orthologue in *A. niger* is important for weak acid resistance

To determine whether orthologues of WarA are present in other species of fungi, a BLAST-search using the *Afu*WarA protein sequence as a query was conducted. In *A. niger,* this identified An08g08340, a protein with 46.2% identity and 63% similarity to the *A. fumigatus* WarA protein. Orthologues of *AfuwarA* were also found to be present in *Penicillium* and *Botrytis spp*.

*Aspergillus niger* is highly resistant to weak acids (especially sorbic and benzoic acids), and can readily cause food and drink spoilage. To determine whether *warA* is also required for weak acid resistance in *A. niger*, *An08g08340* was deleted by a targeted gene replacement approach, with successful deletion confirmed by PCR and Southern Blotting (Fig. S2). Sensitivity of the Δ*An08g08340* deletion strain to different weak acids was evaluated (Fig. 3). In contrast to the ΔwarA deletant of *A. fumigatus*, the *A. niger* Δ*An08g08340* strain demonstrated only slight sensitivity to sorbic acid, which was most apparent when individual conidia of the Δ*warA* strain were spread onto medium containing sorbic acid (Fig. S3A). However, the deletant was highly sensitive to propionic, butanoic and benzoic acids as evaluated by radial growth on agar (Fig. 3). Determination of Minimum Inhibitory Concentrations (MIC) in broth also corroborated the radial growth data; the MIC in benzoic acid was 4.5 ± 0.5 mM in the wild type (WT; n=3), compared with 3.2 ± 0.2 mM in Δ*warA* (n=3). The MIC in butanoic acid was 8.6 ± 0 mM in the WT (n=2), and 6.9 ± 0.3 mM in Δ*warA* (n=3). Resistance was restored to WT levels when *An08g08340* was reintroduced into the Δ*An08g08340* strain (Fig. S4). Thus, *An08g08340* has an important role in weak acid resistance in *A. niger*, and was named as *warA* (Weak Acid Resistance A) also in this species.

**FIG 3.**
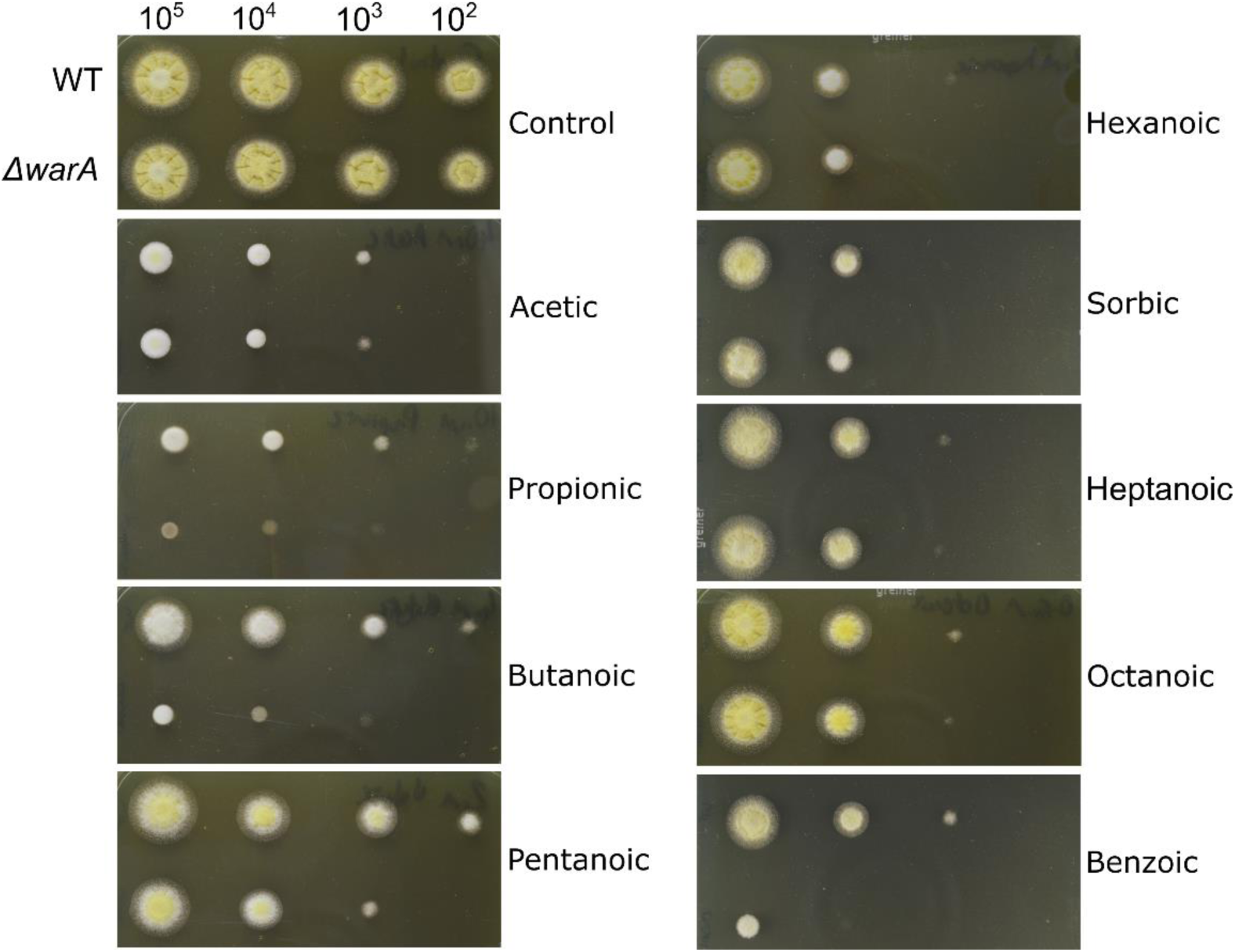
Radial growth of *A. niger* Δ*warA* growing on different weak acids. Plates were inoculated with a 10-fold dilution series of conidial suspensions; approximate numbers of conidia are indicated above the pictures. Images were captured after 2 d growth at 28°C, and are representative of 2-3 independent experiments. Concentrations of acids used are given in the Materials and Methods.

Sorbic acid (and structurally related acids) are known to be detoxified by decarboxylation in *A. niger*, but not in *A. fumigatus* (14, 15). The decarboxylation process involves three linked genes: *cdcA*, *padA* and *sdrA* (12, 14). CdcA is the key enzyme involved in the decarboxylation, whereas SdrA is a transcription factor regulating expression of CdcA and PadA synthesizes a cofactor for CdcA. It was hypothesized that mild sorbic acid sensitivity of *A. niger* Δ*warA* may be due to downregulation of *cdcA*, *padA* or *sdrA* genes in the Δ*warA* strain. To investigate the relationship between WarA and weak acid decarboxylation, a ΔΔ*cdcA/warA* mutant was constructed. The ΔΔ*cdcA/warA* strain was more sensitive to sorbic acid than the Δ*cdcA* strain (Fig. S3B), suggesting a *cdcA*-independent role for *warA* in resistance of *A. niger* to sorbic acid.

### Determination of WarA-regulated genes by transcriptomic analysis during weak acid stress of *A. niger*

The weak acid sensitivity of the Δ*warA* mutant suggested that WarA regulates genes which are important for weak acid resistance. Previously, genes upregulated by sorbic acid exposure in *A. niger* were successfully identified by exposing conidia of the wild-type (WT) to sorbic acid during germination (6). In order to identify which genes are differentially regulated in *A. niger* Δ*warA*, RNA-seq analysis was conducted with WT and Δ*warA* conidia germinated in the presence or absence of sorbic acid. Germination of WT conidia for 1 h in the presence of 1 mM sorbic acid resulted in 3,274 differentially expressed genes (FDR adjusted p-value < 0.05) (1885 upregulated, 1467 downregulated) in comparison with conidia germinated in the absence of sorbic acid. In Δ*warA* conidia, 3,442 genes were differentially expressed during germination in the presence of sorbic acid (1885 upregulated, 1,557 down regulated), in comparison with germination without sorbic acid. Importantly, a number of genes were identified that were highly upregulated in the WT during sorbic acid exposure, but not in Δ*warA* (Table 1 and Table S1). This included a gene encoding benzoate *para*-hydroxylase (*bphA*), an enzyme known to be required for benzoate detoxification (9), which had a Log Fold Change (Log_2_FC) of 6.50 in the WT, compared with a Log_2_FC of −0.51 in Δ*warA.* A number of uncharacterized enzymes also required *warA* for normal upregulation by sorbic acid, e.g., An12g09130 encoding a putative dienelactone hydrolase, and An12g02790 a putative isoflavone reductase (Log_2_FC 5.95 in WT, Log_2_FC 0.36 in Δ*warA*), as well as several genes encoding putative transporter proteins. Of particular interest amongst these transporters was An14g03570, an ABC-type transporter with 56% amino acid sequence similarity to *S. cerevisiae* Pdr12p. Pdr12p has a crucial role in weak acid detoxification in *S. cerevisiae* (17, 18). To support the RNA-seq data, five of the genes showing differential expression between the WT and Δ*warA* strains were selected for qRT-PCR analysis (Fig. 4). The qRT-PCR data supported the trends in gene expression seen in the RNA-seq dataset; all the selected genes had a lower transcript abundance in Δ*warA*, compared with the WT during sorbic acid treatment. In addition, transcript abundances of the selected genes were compared by qRT-PCR during benzoic acid treatment. As expected, all the genes upregulated by sorbic acid were also upregulated by benzoic acid, and had lower transcript abundances in the Δ*warA* mutant, compared with the WT (Fig. 4).

**FIG 4.**
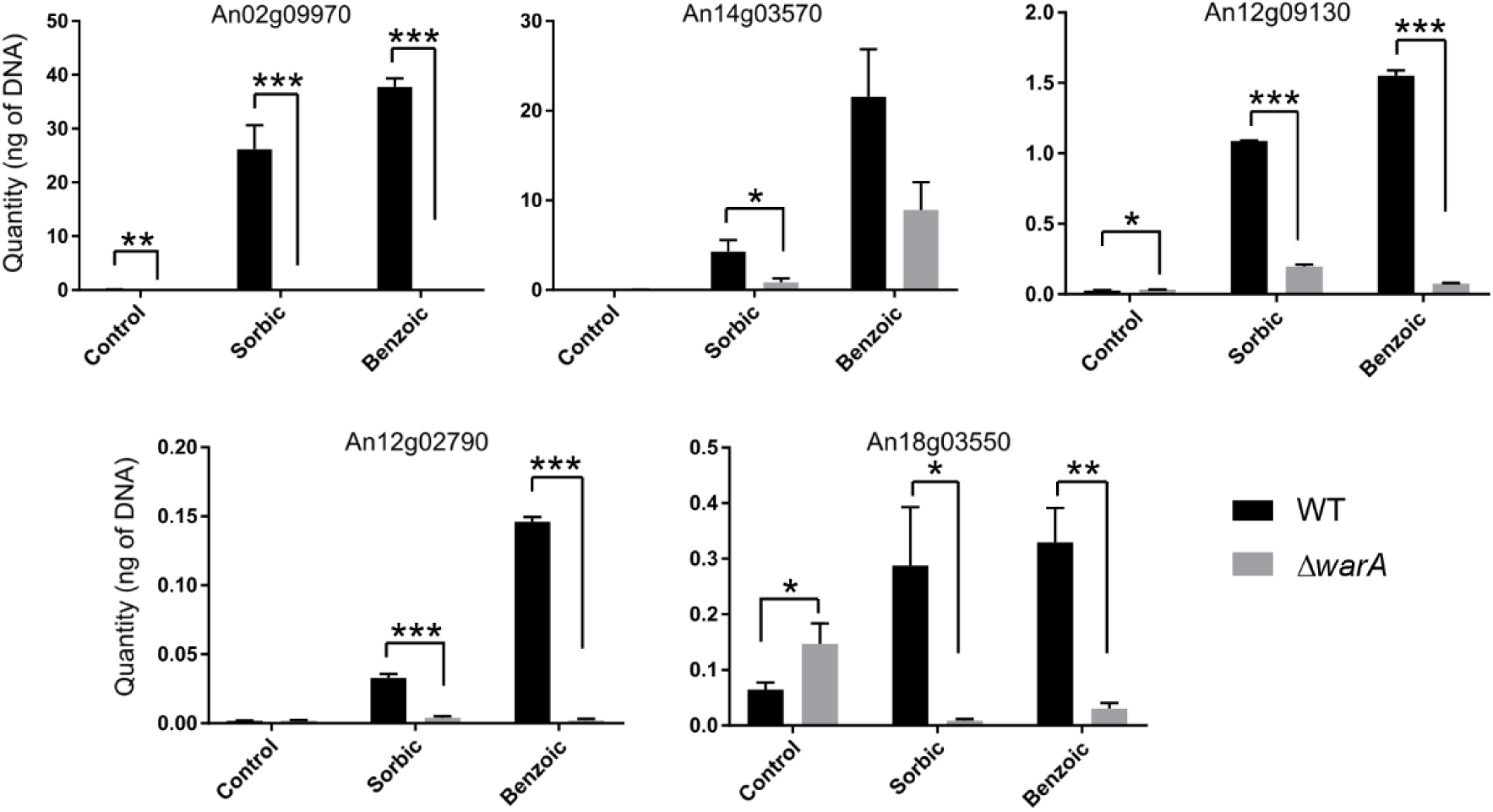
qRT-PCR of genes differentially regulated in WT and Δ*warA* strains of *A. niger.* Transcript abundances in WT (black bars) and Δ*warA* (grey bars) conidia germinated in control media, or in the presence of 1 mM sorbic acid or 1 mM benzoic acid. Error bars are standard deviation of 3 technical replicates. WT and Δ*warA* transcript abundances were compared by Student’s t-test (*p < 0.05, **p <0.01, *** p<0.001).

**TABLE 1.** Transcriptomics data for selected genes upregulated in the WT during sorbic acid treatment and differentially expressed in WT versus Δ*warA*.^a^

| Gene_ID | Log<sub>2</sub>FC<sup>b</sup>.WT<br>_Sorbic vs<br>WT control | Log<sub>2</sub>FC.$\Delta warA$<br>_Sorbic vs<br>$\Delta warA$ control | WT<br>sorbic<br>RPKM<sup>c</sup> | $\Delta warA$<br>sorbic<br>RPKM | Function |
| --- | --- | --- | --- | --- | --- |
| An14g03570 | 9.35 | 7.1 | 611.3 | 146 | ABC-type transporter with similarity to *S.cerevisiae* Pdr12p |
| An12g09130 | 9.31 | 5.98 | 6055.3 | 224.7 | Possible diene lactone hydrolase function |
| An02g09970 | 8.56 | 4.35 | 1532.7 | 7.3 | Ortholog(s) have role in drug response, hexose transport, pathogenesis |
| An13g02460 | 8.24 | 3.08 | 283 | 0.3 | Protein similar to nonribosomal peptide synthases (NRPS-like) |
| An06g02170 | 7.98 | 1.51 | 80.3 | 3 | Ortholog(s) have S-adenosylMet-dependent methyltransferase activity |
| An13g03170 | 7.25 | 5.73 | 1433.3 | 465.3 | Unknown |
| An12g09120 | 7.14 | 2.87 | 312.3 | 31.7 | Unknown |
| An09g03500 | 6.5 | -0.51 | 226.3 | 4 | Putative benzoate-para-hydroxylase; 3-hydroxybenzoate 4- hydroxylase |
| An12g02790 | 5.95 | 0.36 | 135.3 | 2.7 | Isoflavone reductase - phenylcoumaran benzylic ether reductases type |
| An08g07850 | 5.81 | 3.77 | 11809.3 | 3248.7 | Unknown |
| An01g05850 | 5.59 | 1.5 | 75.7 | 3 | Thioesterase domain protein |
| An08g01560 | 5.22 | -1.47 | 802.7 | 5.7 | Ortholog(s) have role in meiotic cell cycle, regulation of TORC1 signaling |
| An04g05240 | 5.02 | 2.84 | 833 | 259.7 | Unknown |
| An11g04385 | 4.9 | 2.02 | 74 | 5.7 | Possible ubiquitin hydrolase |
| An13g02450 | 4.21 | 0 | 492.7 | 0 | Six hairpin glycosidase |
| An08g01980 | 3.31 | 1.71 | 309 | 74.7 | Unknown |
| An13g02290 | 3.06 | -0.95 | 73.3 | 1.7 | Possible 3-dehydroshikimate dehydratase |
| An09g05760 | 2.74 | -0.47 | 397.7 | 40 | Ortholog(s) have actomyosin contractile ring, intermediate layer localization |
| An18g03550 | 2.45 | -2.43 | 1506.7 | 22.3 | Similar to yeast Arr3 arsenate transporter |
| An08g05750 | 2.26 | 0.76 | 1095 | 298 | Unknown |
| An16g00700 | 0.96 | -3.07 | 345 | 20.7 | Has domain(s) with predicted 2Fe,2S cluster binding, oxidoreductase activity |

684 <sup>a</sup>See supplementary Table 1 for full list of genes. <sup>b</sup>Log<sub>2</sub> Fold Change. <sup>c</sup>Reads per Kilobase per Million.

### Characterization of *An02g09970* and *An14g03570*

The transcriptomic analysis identified genes that are downregulated in *A. niger* Δ*warA*, relative to the WT strain, during weak acid stress. These genes may therefore have a role in weak acid resistance. To investigate this, two genes of interest (*An02g09970* and *An14g03570*) were selected for further characterization. *An02g09970* encodes a putative transmembrane transporter of the Major Facilitator Superfamily (MFS), and was selected for further investigation due to its extremely high transcript abundance during sorbic acid treatment, and large disparity in transcript abundance between the WT and Δ*warA* strains (Log_2_FC 8.56 in WT vs. Log_2_FC 4.35 in Δ*warA*) (Table 1). The protein also shares significant sequence similarity with Tpo2 and Tpo3, *S. cerevisiae* proteins involved in resistance to acetic, propionic and benzoic acids (20). *An14g03570* encodes an ABC-type transporter with similarity to the weak acid detoxification protein Pdr12p in *S. cerevisiae*, as stated above. Both genes were deleted in *A. niger* by a targeted gene replacement approach, and mutant genotypes confirmed by PCR and Southern blotting (Fig. S5). Sensitivity of the constructed deletion strains to weak acids was then evaluated. The Δ*An02g09970* mutant did not exhibit altered sensitivity to any of the weak acids tested (Fig. 5). However, the Δ*An14g03570* mutant was more sensitive to sorbic, pentanoic and benzoic acids (Fig. 5 and Table 2), and resistance was restored to WT levels when *An14g03570* was reintroduced into the Δ*An14g03570* strain (Fig. S6). Because of the similarity in sequence and function between An14g03570 and Pdr12p, An14g03570 was named PdrA.

**FIG 5.**
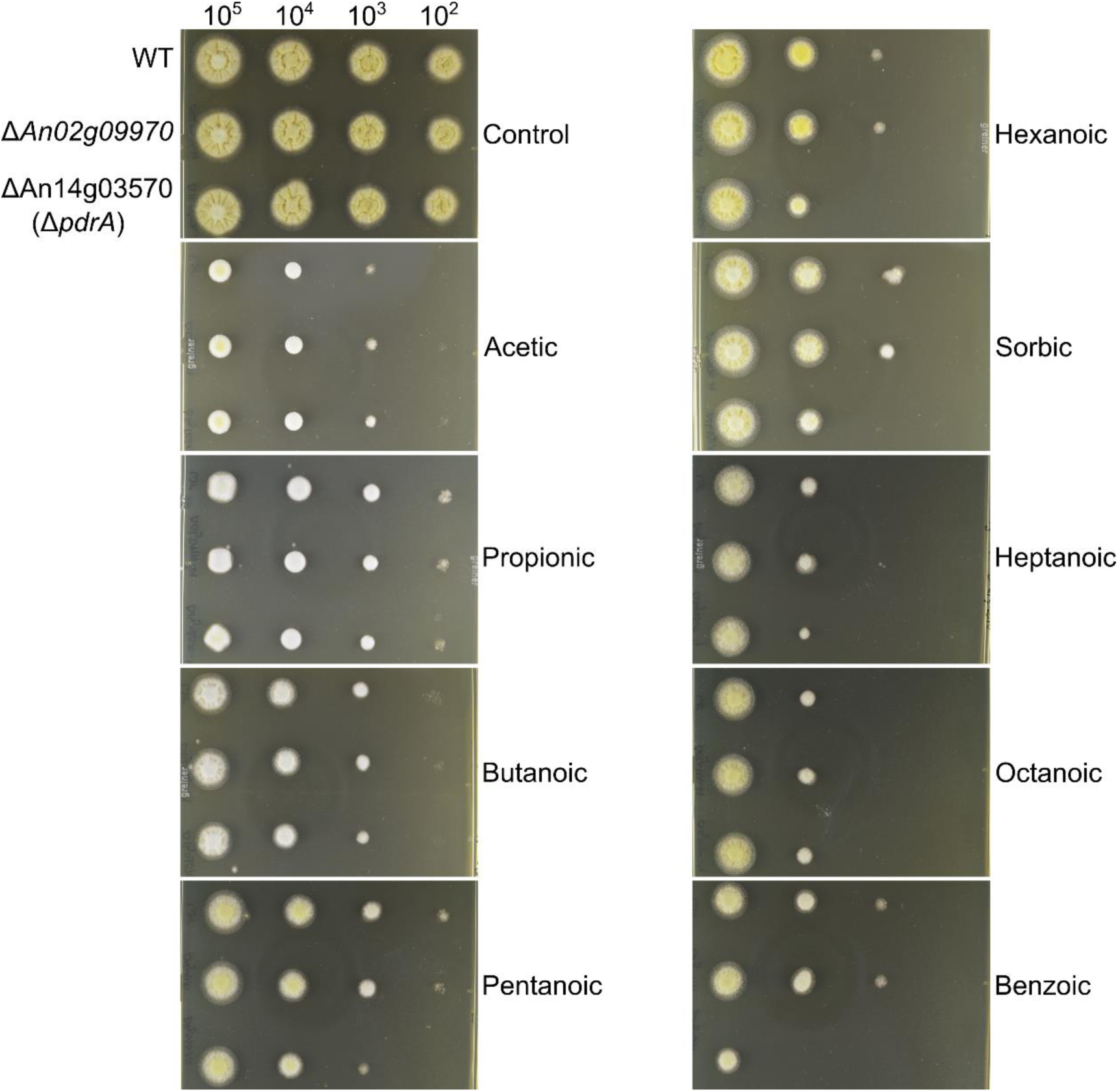
Radial growth of Δ*An02g09970* and Δ*An14g03570 (ΔpdrA)* deletion strains growing on weak acids. Plates were inoculated with a 10-fold dilution series of conidial suspensions.

**TABLE 2.** MIC values (in mM) for *A. niger* WT and the Δ*An14g03570* (Δ*pdrA*) deletion strain.^a^

| Acid | MIC (mM) |  |
| --- | --- | --- |
| | WT | $\Delta pdrA$ |
| Benzoic | 4.67 ± 0.12 | 3.40 ± 0.20 |
| Pentanoic | 3.70 ± 0.35 | 2.60 ± 0.30 |
| Sorbic | 5.00 ± 0.35 | 3.80 ± 0.35 |

<sup>a</sup>Values are averages of 3 biological replicates ± StdDev.

### Complementation of *S. cerevisiae* Δ*pdr12* strain with PdrA (An14g03570)

The above results showed that *pdrA* is required for resistance of *A. niger* to certain weak acids. Because PdrA has significant protein sequence similarity with *S. cerevisiae* Pdr12p (36% sequence identity, 56% similarity), it was hypothesized that these proteins could be functional homologues. To test this hypothesis, functional complementation of the *S. cerevisiae* Δ*pdr12* strain was attempted. The cDNA sequence of *A. niger pdrA* was cloned between the *S. cerevisiae PDR12* promoter and terminator to allow for native regulation of *pdrA* in response to weak acid stress in *S. cerevisiae*. The resulting plasmid (Fig. S7) was transformed into a *S. cerevisiae* Δ*pdr12* deletion strain. Transformants were tested for sensitivity to a range of weak acids. As hypothesized, *pdrA* could indeed functionally complement *PDR12*: sensitivity of the Δ*pdr12* strain to weak acids was largely rescued in cells transformed to express *pdrA* (Fig. 6). The resultant level of resistance was similar to that evident in Δ*pdr12* cells expressing *PDR12* from the same vector backbone.

**FIG 6.**
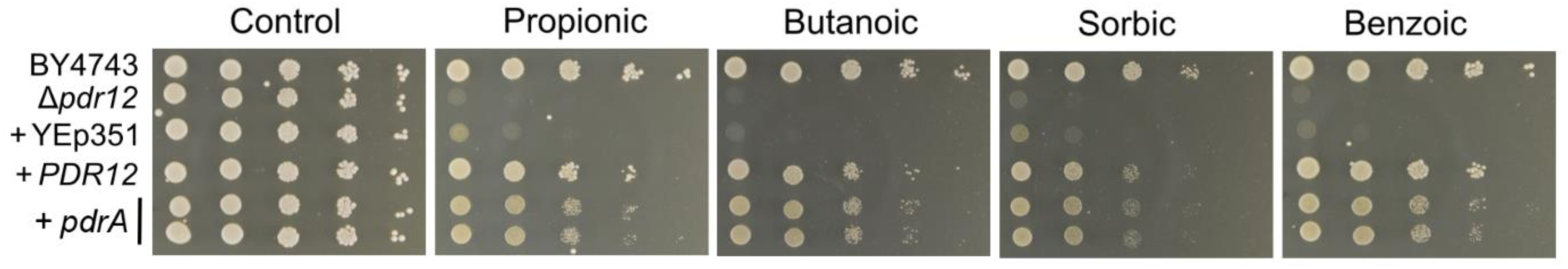
Growth of *S. cerevisiae* complemented strains on weak acids. Ten-fold dilution series of *S. cerevisiae* strains (isogenic with the BY4743 wild type) were inoculated onto medium containing weak acids. The Δ*pdr12* strain was transformed with either empty plasmid (+YEp351), YEp351 plasmid containing *PDR12* (+*PDR12)* or YEp351 plasmid containing the *pdrA* ORF and *PDR12* promoter and terminator (+*pdrA*). Two independent transformants of the +*pdrA* strain are shown.

### *pdrA* is a determinant of heteroresistance to sorbic acid in *A. niger* conidia

Resistance to weak acid preservatives has been found to be heterogeneous between individual cells of genetically-uniform cell populations of the yeasts *Zygosaccharomyces bailii* and *S. cerevisiae* (8, 26, 27). We observed a similar phenomenon in populations of *A. niger* conidia, whereby small subpopulations were capable of germinating and forming colonies at high concentrations (>4 mM) of sorbic acid (Fig. 7A). Conidia harvested from colonies that grew on these high concentrations of sorbic acid did not retain increased resistance upon direct re-inoculation to sorbic acid-containing medium (data not shown), suggesting that these were transient, non-heritable phenotypes, i.e., not due to genotypic variants within the population. To test whether this heteroresistance had its origin in the ungerminated conidial state, conidia were also pre-germinated for 6 hr, before spread plating onto sorbic acid-containing medium. This showed that germinated conidia were much more susceptible to sorbic acid (Fig. 7A). Moreover, resistance to sorbic acid among pre-germinated conidia was much more homogeneous than when the resistance-assay commenced with ungerminated conidia, as evidenced by the gradients of the dose inhibition curves: such dose response curves reflect heterogeneity, with shallower curves indicating greater heterogeneity (23, 33). Thus, at least some factors determining heteroresistance to sorbic acid are specific to ungerminated conidia, and are lost upon germination. Given the contributions of *warA* and *pdrA* to sorbic acid resistance in *A. niger*, it was tested whether these genes could be determinants of heteroresistance. Dose response curves of Δ*pdrA* and Δ*warA* conidia demonstrated that *pdrA* makes a significant contribution to sorbic acid heteroresistance in *A. niger* conidia, whereas *warA* does not (Fig. 7B, data for Δ*warA* not shown).

**FIG 7.**
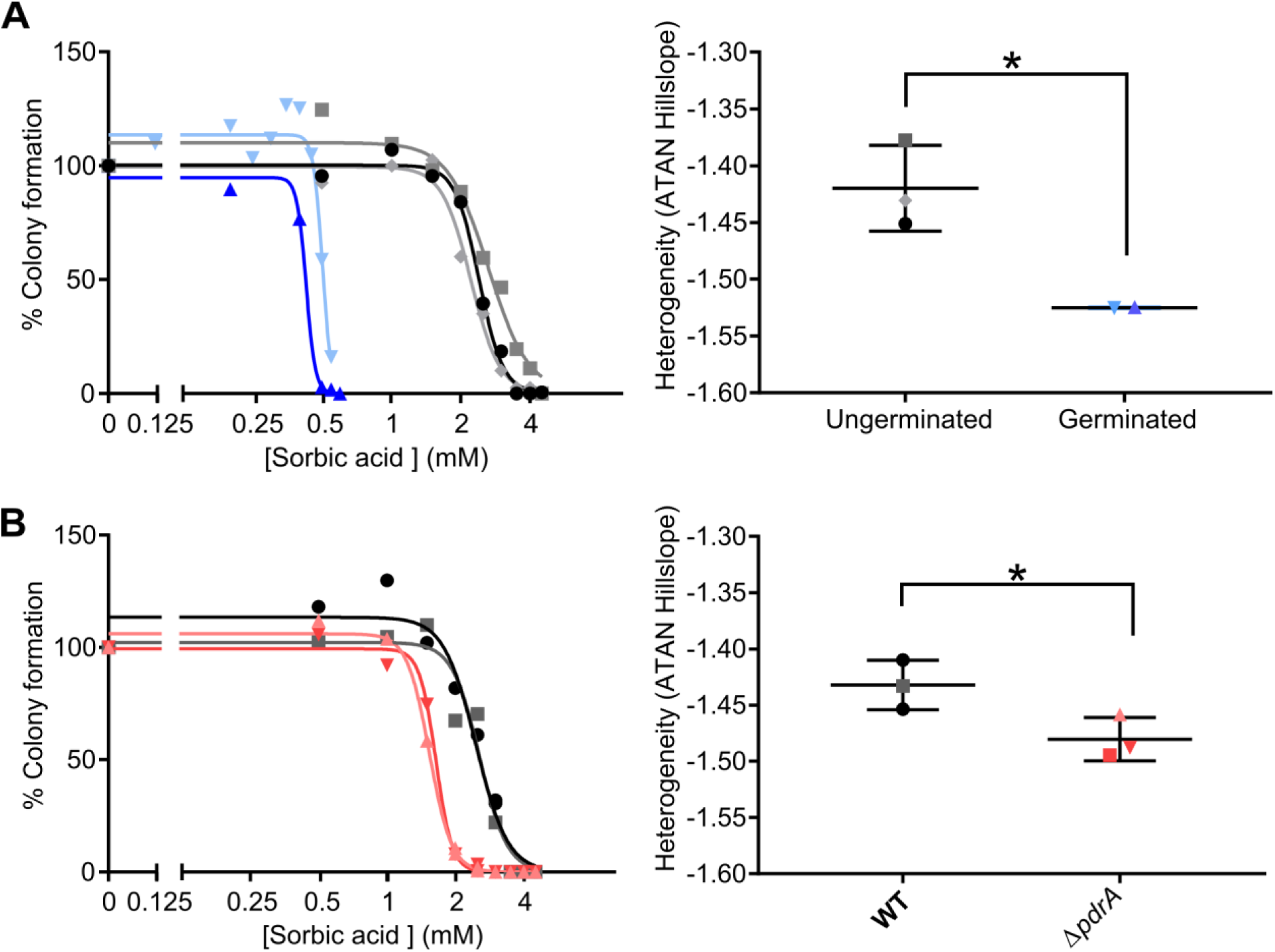
Sorbic acid dose response curves for *A. niger* conidia. (A) Dose response curves of germinated (blue lines) and ungerminated (black/grey lines) WT conidia, and comparison of slope values. Dose response curve slope values were compared with 2-way Welch’s T-test (p= 0.0404) n = 2-3. (B) Dose response curves of WT (black/grey lines) and Δ*pdrA* (pink/red lines) conidia and comparison of dose response curve slope values, compared with 2-way Welch’s T-test (p = 0.0468) n = 3. Two representative independent experiments are shown in the dose response curve.

## DISCUSSION

This study reports the discovery of novel factors determining weak acid resistance in moulds. By screening >400 transcription factor deletion strains in *A. fumigatus*, we discovered a previously uncharacterized transcription factor that is required for resistance to certain weak organic acids. This transcription factor, here named *Weak Acid Resistance A* (*warA*) was also found to be present in the food spoilage mould *A. niger*, where it also plays an important role in weak acid resistance. However, WarA appears to mediate resistance to different weak acids in *A. fumigatus* and *A. niger*. In *A. fumigatus*, the Δ*warA* strain is particularly sensitive to linear-chain acids 3-6 carbons in length, whereas in *A. niger* the Δ*warA* strain is most sensitive to propionic, butanoic and benzoic acids, but exhibited less sensitivity to 5 and 6 carbon acids. Such differences in acid sensitivity may reflect differences in the WarA regulon between *A. niger* and *A. fumigatus*, or the divergence in gene function within the WarA regulon.

We sought to gain insight to the WarA regulon in *A. niger* by conducting a comparative transcriptomics experiment between germinating WT and Δ*warA* conidia treated with sorbic acid. This approach identified several genes that appear to be regulated (either directly or indirectly) by WarA. These include a number of putative enzymes and transporter proteins, offering several candidates for future studies of weak acid resistance mechanisms in *A. niger*. Amongst these candidates, we attempted to characterize two putative transporter-protein genes. The first of these functions, An02g09970, is a transporter of the Major Facilitator Superfamily, with sequence similarity to Tpo2p and Tpo3p in *S. cerevisiae*. Tpo2p and Tpo3p are transporters of the DHA1 (Drug:H+ antiporter-1) family and are known to be required for resistance to acetic, propionic and benzoic acids (20). However, deletion of *An02g09970* did not sensitize *A. niger* to any of the acids tested, and so the role of this gene remains unknown. It is possible that this transporter is responsible for detoxification of other xenobiotics not tested here (if indeed it has a role in detoxification at all), or that *An02g09970* is functionally redundant with other *A. niger* genes. We also attempted to characterize *pdrA*, encoding an ABC-type transporter. Deletion of *pdrA* resulted in increased sensitivity to pentanoic, hexanoic, sorbic and benzoic acids, substantiating a role for this protein in weak acid resistance. Importantly, we were able to demonstrate that *pdrA* is a functional homologue of *PDR12* in *S. cerevisiae*. Pdr12p is a key protein involved in weak acid resistance of *S. cerevisiae* (17), where it is thought to efflux weak acid anions from the cytoplasm in an energy-dependant manner (18). The identification of PdrA as a functional homologue of Pdr12p in a mould species such as *A. niger* shows that a similar mechanism of weak acid detoxification by active efflux may operate in yeasts and moulds. Interestingly, the Δ*pdr12* functional complementation experiment demonstrated that *pdrA* confers resistance to a broader range of weak acids than was suggested by the weak acid sensitivity of the Δ*pdrA* strain. For example, *pdrA* complemented the propionic acid sensitivity of *S. cerevisiae* Δ*pdr12*, but the *A. niger* Δ*pdrA* strain was not more sensitive to propionic acid than the WT. This may indicate the presence of multiple, redundant mechanisms for resistance to certain weak acids in *A.niger*, which may not operate in *S. cerevisiae*.

*pdrA,* as well as several other candidate WarA-regulated genes were all upregulated in response to both sorbic and benzoic acids. This suggests a degree of overlap between transcriptomic responses to different weak acids, as also found in *S. cerevisiae* (34). Thus, although we characterized the WarA regulon by comparative transcriptomics in response only to sorbic acid in the present study, it is likely that many of the differentially expressed genes would be similarly regulated in response to other weak acids. There may be relevant consensus sequences within WarA-regulated genes, although these are not apparent from promoter sequence alignments we have carried out. In *S. cerevisiae*, a *cis*-acting weak acid response element (WARE) was discovered in the promoter of *PDR12* which is required for *PDR12* induction by the transcription factor War1p (19).

The regulation of WarA itself is also an outstanding question. Recent evidence in *S. cerevisiae* suggests that weak acid anions bind directly to the transcription factors War1p and Haa1p, thereby regulating their DNA-binding transcriptional activation (35). However, WarA shares very little sequence homology with either War1p or Haa1p. In fact, a BLAST search of the *S. cerevisiae* protein database with the WarA protein sequence yields no hits at all. Nevertheless, a similar mechanism of transcription factor activation cannot be ruled out for WarA, particularly as direct ligand binding has been established for a number of Zn2Cys6 family transcription factors (of which WarA is a member), including Pdr1p, Pdr3p, Leu3p and Put3p (36–38).

During the course of this study, experiments with sorbic acid determined that genetically-uniform populations of *A. niger* conidia demonstrate heteroresistance to this weak acid. Phenotypic heterogeneity within microbial cell populations has been demonstrated in a number of fungi in response to environmental stresses [reviewed in (23)], however this is the first report of weak acid heteroresistance in fungal conidia. Interestingly, heteroresistance was decreased within 6 h of conidial germination, suggesting that at least some factors underlying this heterogeneity are limited to ungerminated conidia and are lost upon germination. Resistance to sorbic acid was also markedly lower in germinated conidia, which has also recently found to be the case for propionic acid (39). Heteroresistance to weak acids in fungal conidia has significant implications for the food industry, because spoilage of products may occur due to contamination with just a few conidia from a highly resistant subpopulation. Thus, future spoilage control strategies may have to take into account the presence of weak acid heteroresistance, perhaps by specifically targeting resistant subpopulations.

Conidia of the Δ*pdrA* strain showed a significantly more homogeneous response to sorbic acid. Heteroresistance typically arises from gene expression heterogeneity (or noise) (40), so the present results suggest that *pdrA* could be expressed heterogeneously within conidial populations; the conidia expressing more *pdrA* potentially able to withstand sorbic acid stress. It is also noted that deletion of *pdrA* did not eliminate sorbic acid heteroresistance in *A. niger*, so it likely that other genes also contribute.

In summary, this study markedly advances our understanding of weak acid resistance mechanisms in *A. niger*. The identification of WarA as a key transcription factor involved in weak acid resistance allowed us in turn to identify many more genes which may also be important. Further work is required to determine how all these genes may contribute to weak acid resistance. Moreover, we demonstrated here that a key weak acid resistance mechanism operates in both *S. cerevisiae* and *A. niger*, in the form of the functionally homologous ABC-transporters Pdr12p and PdrA respectively.

## MATERIALS AND METHODS

### Strains and media

The *Aspergillus fumigatus* transcription factor deletant library, derived from wild-type strain MFIG001, was constructed by homologous recombination using gene replacement cassettes and transformation methodologies as described (30, 41). Studies in *Aspergillus niger* were performed in the *A.niger* N402 background (referred to as the *A. niger* WT throughout) and an *A. niger ΔcdcA* deletion strain (14). *Aspergillus* strains were cultivated on slopes of Potato Dextrose Agar (PDA) (Sigma) for 7 days at 28°C. Conidia were harvested using 0.1% (v/v) Tween 80 and filtered through a 40 μm cell strainer (Fisher), before counting on a haemocytometer. Studies in *S. cerevisiae* used the BY4743 background and isogenic *Δpdr12* deletion strain, cultivated on YEPD agar (2% glucose, 2% bactopeptone (Oxoid), 1% yeast extract (Oxoid), 1.5% agar) at 30°C. The *S. cerevisiae* strains were obtained from EUROSCARF (Frankfurt). Growth assays with weak acids (below) were performed on YEPD agar (pH 4).

### Deletant-library screening and growth assays

The first round of *A. fumigatus* transcription factor deletion library screening was performed in a 96-well array format. Conidial suspensions of the *A. fumigatus* strains were initially supplied in 40% glycerol-0.01% PBS solution at a concentration of 4 x 10^7^ ml^-1^. These were subsequently arrayed in 96-well plates at a concentration of 4 x 10^5^ conidia ml^-1^ in 0.01% Tween 20, and transferred using a 96-pin tool, to Nunc™ Omnitray™ single-well plates containing YEPD agar (pH 4), then incubated at 28°C for 2 - 3 days. Radial growth was measured using ImageJ, and compared between control medium, and medium containing sorbic acid. The second round of screening was performed on 90 mm Petri dishes. Plates were inoculated with 10^5^ conidia, and incubated at 37°C for 3 days. Radial growth was compared between control medium (YEPD) and the same medium containing sorbic acid.

Subsequent growth assays with weak acids on solid medium were performed in *Aspergillus* spp. by inoculating YEPD agar (pH 4) supplemented with weak acids, with 5 μl of conidial suspension, containing 10^5^-10^2^ conidia and subsequent incubation at 28°C for 2 – 3 days. Weak acid concentrations used for *A. fumigatus* were: 15 mM acetic acid, 4 mM propionic acid, 1.5 mM butanoic acid, 0.75 mM pentanoic acid, 0.2 mM sorbic acid, 0.25 mM hexanoic acid, 0.5 mM benzoic acid, 0.08 mM heptanoic acid, 0.05 mM octanoic acid. Weak acid concentrations used for *A. niger*: 40 mM acetic acid, 10 mM propionic acid, 4 mM butanoic acid, 2 mM pentanoic acid, 1.5 mM sorbic acid, 2 mM hexanoic acid, 2 mM benzoic acid, 1 mM heptanoic acid, 0.75 mM octanoic acid.

Minimum inhibitory concentrations (MICs) of weak acids were determined by placing 10 ml of YEPD broth (pH 4) into 30 ml McCartney bottles, and inoculating with 10^4^ conidia. Bottles were incubated statically at 28°C for 28 days, and the concentration of acids required to completely inhibit visible growth recorded. Concentrations of acids used were at 0.2 mM increments for benzoic and sorbic acids, 0.3 mM increments for pentanoic and butanoic acids, and 2 mM increments for propionic acid.

Dose response curves were generated by harvesting conidia as stated above, diluting to 500 spores ml^-1^ and spreading 200 μl of this onto YEPD agar (pH 4) containing sorbic acid. Plates were incubated at 28°C for up to 28 days and colonies counted. For pre-germinated conidia, conidia were first inoculated into 10 ml of YEPD/Tween 80 (6.66 ml YEPD/3.33 ml 0.1 % Tween 80) to a final concentration of 500 spores ml^-1^ and incubated statically at 28°C for 6 hr, before spread plating and incubation as above. For quantitative comparison of heteroresistance, Hill slopes were fitted to plots (% Viability vs. log10[sorbic acid]) using Prism software and arctangent values for the slopes calculated with Excel to estimate relative heterogeneity (a shallower slope indicating higher heterogeneity) (33, 42).

### RNAseq and qRT-PCR

For preparation of RNA, conidia of *A. niger* N402 were inoculated into 1 litre of YEPD broth (pH 4) to a final concentration of 10^6^ conidia ml^-1^ and incubated at 28°C for 1 hr, with shaking at 150 rev. min^-1^. For sorbic acid or benzoic acid treatments, the medium was supplemented with 1 mM sorbic or 1 mM benzoic acid for the 1 h incubation: these concentrations inhibit conidial germination over the course of the experiment, but are not lethal (∼25% of the MIC values for these acids). Conidia were harvested by filtration through a Corning vacuum filtration unit, and immediately used for RNA extraction. RNA was extracted using a Norgen Biotek Plant/Fungi Total RNA extraction kit, as per the manufacturer’s instructions.

RNAseq analysis was performed by the University of Liverpool Centre for Genomic Research. Three biological replicates were performed for each timepoint in each condition. Between 463-1000ng of total RNA (depending on available material) was Poly A-treated using the NEBNext® Poly(A) mRNA Magnetic Isolation Module, and subsequently purified using Ampure RNA XP beads. Successful depletion of rRNA was confirmed using Qubit fluorometric quantification (ThermoFisher) and Agilent 2100 Bioanalyzer. All of the depleted RNA was used as input material for the NEBNext® Ultra™ Directional RNA Library Prep Kit for Illumina®. Following 15 cycles of amplification the libraries were purified using Ampure XP beads. Each library was quantified using Qubit and the size distribution assessed using the Bioanalyzer. These final libraries were pooled in equimolar amounts using the Qubit and Bioanalyzer data. The quantity was assessed using a Qubit® dsDNA HS Assay Kit, while the quality and average fragment size was assessed using the High Sensitivity DNA Kit on the Agilent Bioanalyzer. The RNA libraries were sequenced on an Illumina® HiSeq 4000 platform with version 1 chemistry using sequencing by synthesis (SBS) technology to generate 2 x 150 bp paired-end reads. Initial processing and quality assessment of the sequence data was performed as follows. Briefly, base calling and de-multiplexing of indexed reads was performed by CASAVA version 1.8.2 (Illumina). The raw FASTQ files were trimmed to remove Illumina adapter sequences using Cutadapt version 1.2.1 (43). The option “-O 3” was set, so the 3’ end of any reads which matched the adapter sequence over a stretch of at least 3 bp was trimmed off. The reads were further trimmed to remove low quality bases, using Sickle version 1.200 with a minimum window quality score of 20. After trimming, reads shorter than 20 bp were removed. Reads were aligned to the *A.niger* CBS 588.13 genome sequence (http://www.aspergillusgenome.org/download/sequence/A_niger_CBS_513_88/current/A_niger_CBS_513_88_current_chromosomes.fasta.gz) using Tophat version 2.1.0 (44). The expression of each gene was calculated from the alignment files using HTseq-count (45). The raw count data were also converted into FPKM (Fragments per Kilobase per Million reads) values. The count numbers per gene were used during the subsequent differential expression analysis. All of the DGE (Differential Gene Expression) analyses were performed in R (version 3.3.3) environment using the DESeq2 package (46). Significantly differentially-expressed genes were defined as those with FDR-adjusted P-value < 0.05.

For qRT-PCR analysis of gene expression, RNA was extracted as stated above. Genomic DNA was removed by Turbo DNAse Free kit (Invitrogen). cDNA was synthesized using Superscript IV reverse transcriptase (Invitrogen) and Oligo d(T)_20_ primer (Invitrogen), according to the manufacturer’s instructions. Transcripts were amplified using SYBR green Master Mix on an Applied Biosystems 7500 Real-Time PCR instrument and quantified against a standard curve of *A. niger* gDNA. Primer pairs used are listed in Table S2.

### Gene deletion studies and complementation in *A. niger*

Gene deletion studies were performed in *A. niger* N402, the Open Reading Frames (ORFs) of the target genes being replaced by a hygromycin resistance cassette (Fig. S2 and S6). Gene deletion cassettes were constructed by Gap-Repair cloning in *S. cerevisiae* (47). Briefly, the hygromycin resistance cassette, and approximately 1 kb upstream and downstream flanking regions of each target gene were amplified from genomic DNA by PCR (Primers listed in Table S2). The hygromycin resistance cassette had a 20-30 bp homology with the 1 kb flanking regions, and each flanking region also had 20-30 bp homology with the multiple cloning site of the YEp351 plasmid. The PCR products and *Hin*dIII-linearized YEp351 plasmid were transformed into *S. cerevisiae* BY4743, and transformants selected by leucine prototrophy. Successful construction of the gene deletion cassettes was confirmed by PCR. The resulting gene deletion cassettes were amplified by PCR and purified using PCR purification columns (Machery-Nagel), to produce a final linear gene deletion cassette. All PCR reactions were performed using Phusion High-Fidelity DNA Polymerase (New England Biolabs). Production of protoplasts and their transformation was performed using standard methods (48). Transformants were selected using 200 μg ml^-1^ hygromycin (Roche) and confirmed by PCR and Southern blotting (Fig. S2 and S6), using standard methods (49).

For complementation of *A. niger* gene-deletion strains, the genes in question (*warA* and *pdrA* (*An14g03570*)) were amplified by PCR (primers listed in Table S2) and cloned into the *Sbf*I site of the pAN7.1*BAR* plasmid (Fig. S8), which contains the *BAR* gene as a selectable marker (replacing the original hygromycin resistance cassette (50), and imparting resistance to phosphinothricin). PCR amplification included ∼1 kb upstream and ∼300bp downstream of the ORF. Transformation of the resulting plasmids was performed as described above, except that transformants were selected using 5 mg ml^-1^ DL-phosphinothricin (Carbosynth) in YDA agar (Yeast nitrogen base without amino acids, including 1.7 g l^-1^ ammonium sulfate, 10 g l^1^ glucose, 2.25 g l^-1^ ammonium nitrate and 1 M sucrose; pH adjusted to 7.0 using Na_2_HPO_4_, solidified with 1.2% (w/v) agar) (51). Transformants were subjected to an additional round of selection by growth on YDA agar containing 5 mg ml^-1^ DL-phosphinothricin.

### Cloning and complementation in *S. cerevisiae*

Complementation studies were performed in the *S. cerevisiae* Δ*pdr12* strain. The complementation plasmid (Fig. S7) was constructed by yeast gap-repair cloning (47). Briefly the *PDR12* promoter and terminator, and PdrA ORF were amplified by PCR (primers listed in Table S2). Each amplified fragment included a 20-30 bp region of homology either with the YEp351 plasmid or with a neighbouring fragment. The PCR products and *Hin*dIII linearized YEp351 plasmid were transformed into *S. cerevisiae Δpdr12*, and transformants selected by leucine prototrophy (47). Complementation was also performed with the *PDR12* ORF, as a positive control for successful complementation. The *S. cerevisiae Δpdr12* strain was also transformed with the empty YEp351 plasmid, as a negative control.

## ACKNOWLEDGMENTS

This work was supported by the Biotechnology and Biological Sciences Research Council (Grant Number BB/N017129/1). This is an Industry Partnering Award in conjunction with Lucozade Ribena Suntory and Mologic Ltd. This work was also supported by the Wellcome trust (Grant Number 208396/Z/17/Z).

## LEGENDS TO SUPPLEMENTARY FIGURES (data provided in separate attachments)

**FIG S1** Radial growth of the *A. fumigatus* Δ*metR* strain on medium containing 0.5 mM sorbic acid, with or without 0.5 mM methionine.

**FIG S2** PCR and Southern Blot confirmation of *warA* deletion. (A) Targeted gene deletion strategy. The *warA* ORF was replaced with a hygromycin resistance cassette. (B) PCR confirmation of *warA* deletion. Primer pair 1 was used to confirm deletion of *warA* ORF – deletion strains are negative, WT is positive. Primer pair 2 was used to confirm integration of *HygR* at the *warA* locus. (C) Southern blotting of *warA* and *cdcA/warA* deletion strains. gDNA of strains was digested with *Hin*dIII restriction enzyme. Membranes were hybridised with a probe consisting of Digoxigenin-UTP labelled *HygR*. Single bands confirm single integration of deletion cassette into the *A. niger* genome. Transformants Δ*warA_12* and ΔΔ*cdcA/warA_12* were used for experiments.

**FIG S3** (A) Growth of the Δ*warA* strain conidia on medium containing sorbic acid. Plates were inoculated with ∼100 conidia and incubated for 2 d (control medium) or 3 d (medium containing 1 mM sorbic acid). (B) Radial growth of ΔΔ*cdcA/warA* strain on medium containing 1 mM sorbic acid. Plates were inoculated with a 10-fold dilution series of conidial suspensions; approximate numbers of conidia are indicated at the top. Images were captured after 2 d growth at 28°C, and are representative of 2-3 independent experiments.

**FIG S4** Radial growth of complemented Δ*warA* strains on medium containing 2 mM benzoic acid. Plates were inoculated with a 10-fold dilution series of conidial suspensions. Approximate numbers of conidia are indicated above the pictures. Images were captured after 2 d growth at 28°C. Two independent complemented lines are shown.

**FIG S5** PCR and Southern Blotting of *An02g09970* and *An14g03570* (*pdrA*) deletion strains. (A) Targeted gene deletion strategy. The *An02g09970* and *An14g03570* ORFs were replaced with a hygromycin resistance cassette. (B) PCR confirmation of *An02g09970* deletion. Primer pair 1 was used to confirm deletion of the *An02g09970* ORF – deletion strains are negative, WT is positive. Primer pairs 2 and 3 were used to confirm integration of *HygR* at the *An02g09970* locus. (C) PCR confirmation of *An14g03570* deletion. Primer pair 1 was used to confirm deletion of the *An14g03570* ORF – deletion strains are negative, WT is positive. Primer pairs 2 and 3 were used to confirm integration of *HygR* at the *An14g03570* locus. (D) Southern blotting of ΔA*n02g09970* and ΔA*n14g03570* deletion strains. gDNA of strains was digested with the restriction enzymes *Xba*I (for ΔA*n02g09970*) or *Eco*RV (for ΔA*n14g03570*). Membranes were hybridised with a probe consisting of Digoxigenin-UDP labelled *HygR*. Single bands confirm single integration of deletion cassette into the *A. niger* genome (multiple integrations apparent for Δ*An02g09970_39*). Transformants ΔA*n14g03570_4* and Δ*An02g09970_29* were used for experiments.

**FIG S6** Radial growth of complemented Δ*pdrA* strains on medium containing 2mM benzoic acid. Plates were inoculated with a 10-fold dilution series of conidial suspensions. Two independent complemented lines are shown. A Δ*pdrA* strain containing the empty pAN7.1*BAR* plasmid is also shown.

**FIG S7** Plasmid map of YEp351 containing *An14g03570* (*pdrA*)

**FIG S8** Plasmid map of pAN7.1*BAR*.

**TABLE S1** RPKM and Log_2_FC values for *A. niger* genes (Excel file).

**TABLE S2** List of primers used in this study (Excel file).

## REFERENCES

1. Bondi M, Messi P, Halami PM, Papadopoulou C, de Niederhausern S. 2014. Emerging microbial concerns in food safety and new control measures. BioMed Res Int 2014:251512.

2. Kaczmarek M, Avery SV, Singleton I. 2019. Microbes associated with fresh produce: Sources, types and methods to reduce spoilage and contamination. Adv Appl Microbiol 107:29–82.

3. Salmond CV, Kroll RG, Booth IR. 1984. The effect of food preservatives on pH homeostasis in *Escherichia coli*. Microbiology 130:2845–2850.

4. Plumridge A, Hesse SJA, Watson AJ, Lowe KC, Stratford M, Archer DB. 2004. The weak acid preservative sorbic acid inhibits conidial germination and mycelial growth of *Aspergillus niger* through intracellular acidification. Appl Environ Microbiol 70:3506–3511.

5. Freese E, Sheu CW, Galliers E. 1973. Function of lipophilic acids as antimicrobial food additives. Nature 241:321–325.

6. Novodvorska M, Stratford M, Blythe MJ, Wilson R, Beniston RG, Archer DB. 2016. Metabolic activity in dormant conidia of *Aspergillus niger* and developmental changes during conidial outgrowth. Fungal Genet Biol 94:23–31.

7. Warth AD. 1989. Relationships between the resistance of yeasts to acetic, propanoic and benzoic acids and to methyl paraben and pH. Int J Food Microbiol 8:343–349.

8. Stratford M, Steels H, Nebe-von-Caron G, Novodvorska M, Hayer K, Archer DB. 2013. Extreme resistance to weak-acid preservatives in the spoilage yeast *Zygosaccharomyces bailii*. Int J Food Microbiol 166:126–134.

9. Boschloo JG, Moonen E, van Gorcom RFM, Hermes HFM, Bos CJ. 1991. Genetic analysis of *Aspergillus niger* mutants defective in benzoate-4-hydroxylase function. Curr Genet 19:261–264.

10. Fraser JA, Davis MA, Hynes MJ. 2002. The genes *gmdA*, encoding an amidase, and *bzuA*, encoding a cytochrome P450, are required for benzamide utilization in *Aspergillus nidulans*. Fungal Genet Biol 35:135–146.

11. Stratford M, Plumridge A, Pleasants MW, Novodvorska M, Baker-Glenn CAG, Pattenden G, Archer DB. 2012. Mapping the structural requirements of inducers and substrates for decarboxylation of weak acid preservatives by the food spoilage mould *Aspergillus niger*. Int J Food Microbiol 157:375–383.

12. Lubbers RJM, Dilokpimol A, Navarro J, Peng M, Wang M, Lipzen A, Ng V, Grigoriev IV, Visser J, Hilden KS, de Vries RP. 2019. Cinnamic acid and sorbic acid conversion are mediated by the same transcriptional regulator in *Aspergillus niger*. Front Bioeng Biotechnol 7:249.

13. Payne KAP, White MD, Fisher K, Khara B, Bailey SS, Parker D, Rattray NJW, Trivedi DK, Goodacre R, Beveridge R, Barran P, Rigby SEJ, Scrutton NS, Hay S, Leys D. 2015. New cofactor supports α, β-unsaturated acid decarboxylation via 1, 3-dipolar cycloaddition. Nature 522:497–501.

14. Plumridge A, Melin P, Stratford M, Novodvorska M, Shunburne L, Dyer PS, Roubos JA, Menke H, Stark J, Stam H, Archer DB. 2010. The decarboxylation of the weak-acid preservative, sorbic acid, is encoded by linked genes in *Aspergillus spp*. Fungal Genet Biol 47:683–692.

15. Plumridge A, Stratford M, Lowe KC, Archer DB. 2008. The weak-acid preservative sorbic acid is decarboxylated and detoxified by a phenylacrylic acid decarboxylase, PadA1, in the spoilage mold *Aspergillus niger*. Appl Environ Microbiol 74:550–552.

16. Stratford M, Plumridge A, Archer DB. 2007. Decarboxylation of sorbic acid by spoilage yeasts is associated with the *PAD1* gene. Appl Environ Microbiol 73:6534–6542.

17. Piper P, Mahé Y, Thompson S, Pandjaitan R, Holyoak C, Egner R, Mühlbauer M, Coote P, Kuchler K. 1998. The Pdr12 ABC transporter is required for the development of weak organic acid resistance in yeast. EMBO J 17:4257–4265.

18. Holyoak CD, Bracey D, Piper PW, Kuchler K, Coote PJ. 1999. The *Saccharomyces cerevisiae* weak-acid-inducible ABC transporter Pdr12 transports fluorescein and preservative anions from the cytosol by an energy-dependent mechanism. J Bacteriol 181:4644–4652.

19. Kren A, Mamnun YM, Bauer BE, Schuller C, Wolfger H, Hatzixanthis K, Mollapour M, Gregori C, Piper P, Kuchler K. 2003. War1p, a novel transcription factor controlling weak acid stress response in yeast. Mol Cell Biol 23:1775–1785.

20. Fernandes AR, Mira NP, Vargas RC, Canelhas I, Sá-Correia I. 2005. *Saccharomyces cerevisiae* adaptation to weak acids involves the transcription factor Haa1p and Haa1p-regulated genes. Biochem Biophys Res Commun 337:95–103.

21. Mollapour M, Fong D, Balakrishnan K, Harris N, Thompson S, Schüller C, Kuchler K, Piper PW. 2004. Screening the yeast deletant mutant collection for hypersensitivity and hyper-resistance to sorbate, a weak organic acid food preservative. Yeast 21:927–946.

22. Mira NP, Palma M, Guerreiro JF, Sá-Correia I. 2010. Genome-wide identification of *Saccharomyces cerevisiae* genes required for tolerance to acetic acid. Microb Cell Fact 9:79.

23. Hewitt SK, Foster DS, Dyer PS, Avery SV. 2016. Phenotypic heterogeneity in fungi: importance and methodology. Fungal Biol Rev 30:176–184.

24. Band VI, Weiss DS. 2019. Heteroresistance: A cause of unexplained antibiotic treatment failure? PLoS Pathog 15:e1007726.

25. Andersson DI, Nicoloff H, Hjort K. 2019. Mechanisms and clinical relevance of bacterial heteroresistance. Nature Rev Microbiol 17:1.

26. Stratford M, Steels H, Nebe-von-Caron G, Avery SV, Novodvorska M, Archer DB. 2014. Population heterogeneity and dynamics in starter culture and lag phase adaptation of the spoilage yeast *Zygosaccharomyces bailii* to weak acid preservatives. Int J Food Microbiol 181:40–47.

27. Fernández-Niño M, Marquina M, Swinnen S, Rodríguez-Porrata B, Nevoigt E, Ariño J. 2015. The cytosolic pH of individual *Saccharomyces cerevisiae* cells is a key factor in acetic acid tolerance. Appl Environ Microbiol 81:7813–7821.

28. Teertstra WR, Tegelaar M, Dijksterhuis J, Golovina EA, Ohm RA, Wösten HAB. 2017. Maturation of conidia on conidiophores of *Aspergillus niger*. Fungal Genet Biol 98:61–70.

29. Geoghegan IA, Stratford M, Bromley M, Archer DB, Avery SV. 2019. Weak Acid Resistance A (WarA), a novel transcription factor required for regulation of weak-acid resistance and spore-spore heterogeneity in *Aspergillus niger*. BioRxiv https://doi.org/10.1101/788141.

30. Takanori F, van Rhijn, N., Fraczek, M., Gsaller, F., Davies, E., Carr, P., Gago, S., Fortune-Grant, R., Rahman, S., Mabey Glisenan, J., Houlder, E., Kowalski, C.H., Raj, S., Paul, S., Parker, J.E., Kelly, S., Cramer, R.A., Latge, J.-P., Cook, P., Moye-Rowley, S., Bignell, E., Bowyer, P., Bromley, M.J. The Negative Cofactor 2 complex is a master regulator of drug resistance in *Aspergillus fumigatus*. Nature Comm In revision.

31. Natorff R, Sieńko M, Brzywczy J, Paszewski A. 2003. The *Aspergillus nidulans metR* gene encodes a bZIP protein which activates transcription of sulphur metabolism genes. Mol Microbiol 49:1081–1094.

32. Melin P, Stratford M, Plumridge A, Archer DB. 2008. Auxotrophy for uridine increases the sensitivity of *Aspergillus niger* to weak-acid preservatives. Microbiology 154:1251–1257.

33. Holland SL, Reader T, Dyer PS, Avery SV. 2014. Phenotypic heterogeneity is a selected trait in natural yeast populations subject to environmental stress. Environ Microbiol 16:1729–1740.

34. Legras JL, Erny C, Le Jeune C, Lollier M, Adolphe Y, Demuyter C, Delobel P, Blondin B, Karst F. 2010. Activation of two different resistance mechanisms in *Saccharomyces cerevisiae* upon exposure to octanoic and decanoic acids. Appl Environ Microbiol 76:7526–7535.

35. Kim MS, Cho KH, Park KH, Jang J, Hahn JS. 2018. Activation of Haa1 and War1 transcription factors by differential binding of weak acid anions in *Saccharomyces cerevisiae*. Nucleic Acids Res doi:10.1093/nar/gky1188.

36. Thakur JK, Arthanari H, Yang F, Pan S-J, Fan X, Breger J, Frueh DP, Gulshan K, Li DK, Mylonakis E. 2008. A nuclear receptor-like pathway regulating multidrug resistance in fungi. Nature 452:604–609.

37. Sze J-Y, Woontner M, Jaehning JA, Kohlhaw GB. 1992. *In vitro* transcriptional activation by a metabolic intermediate: activation by Leu3 depends on alpha-isopropylmalate. Science 258:1143–1145.

38. Sellick CA, Reece RJ. 2003. Modulation of transcription factor function by an amino acid: activation of Put3p by proline. EMBO J 22:5147–5153.

39. Dijksterhuis J, Meijer M, van Doorn T, Houbraken J, Bruinenberg P. 2019. The preservative propionic acid differentially affects survival of conidia and germ tubes of feed spoilage fungi. Int J Food Microbiol 306:108258.

40. Avery SV. 2006. Microbial cell individuality and the underlying sources of heterogeneity. Nature Rev Microbiol 4:577–587.

41. Zhao C, Fraczek, M. G., Dineen, L., Lebedinec, R., Macheleidt, J., Delneri, D., Bowyer, P., Brakhage, A.A., Bromley, M. 2019. High-throughput gene replacement in *Aspergillus fumigatus*. Curr Protoc Microbiol 54:e88.

42. Stratford M, Steels H, Novodvorska M, Archer DB, Avery SV. 2019. Extreme osmotolerance and halotolerance in food-relevant yeasts and the role of glycerol-dependent cell individuality. Front Microbiol 9:3238.

43. Martin M. 2011. Cutadapt removes adapter sequences from high-throughput sequencing reads. EMBnet J 17:10–12.

44. Langmead B, Salzberg SL. 2012. Fast gapped-read alignment with Bowtie 2. Nat Methods 9:357–359.

45. Anders S, Pyl PT, Huber W. 2015. HTSeq—a Python framework to work with high-throughput sequencing data. Bioinformatics 31:166–169.

46. Love MI, Huber W, Anders S. 2014. Moderated estimation of fold change and dispersion for RNA-seq data with DESeq2. Genome Biol 15:550.

47. Oldenburg KR, Vo KT, Michaelis S, Paddon C. 1997. Recombination-mediated PCR-directed plasmid construction in vivo in yeast. Nucleic Acids Res 25:451–452.

48. Ballance DJ, Turner G. 1985. Development of a high-frequency transforming vector for *Aspergillus nidulans*. Gene 36:321–331.

49. Green MR, Sambrook J. 2012. Molecular Cloning: a Laboratory Manual. Cold Spring Harbor Laboratory Press.

50. Punt PJ, Oliver RP, Dingemanse MA, Pouwels PH, van den Hondel CAMJJ. 1987. Transformation of *Aspergillus* based on the hygromycin B resistance marker from *Escherichia coli*. Gene 56:117–124.

51. Ahuja M, Punekar NS. 2008. Phosphinothricin resistance in *Aspergillus niger* and its utility as a selectable transformation marker. Fungal Genet Biol 45:1103–1110.

