## Supplementary figures for "Weak Acid Resistance A (WarA), a novel transcription factor required for regulation of weak-acid resistance and spore-spore heterogeneity in *Aspergillus niger*"

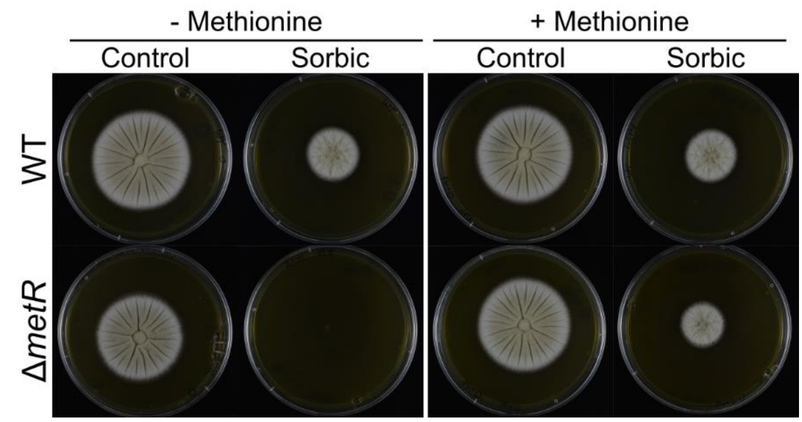


**FIG S1** Radial growth of the *A. fumigatus* Δ*metR* strain on medium containing 0.5 mM sorbic acid, with or without 0.5 mM methionine.

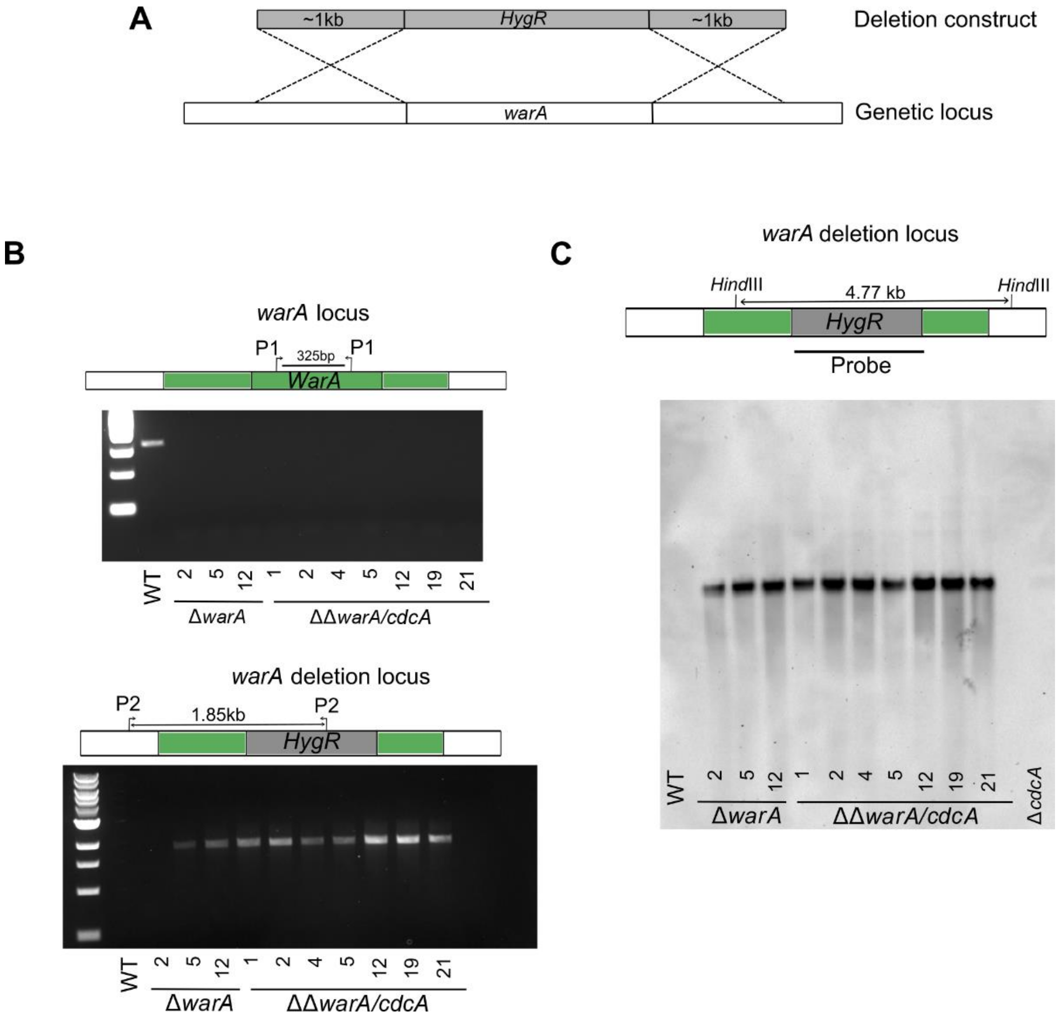


**FIG S2** PCR and Southern Blot confirmation of *warA* deletion. (A) Targeted gene deletion strategy. The *warA* ORF was replaced with a hygromycin resistance cassette. (B) PCR confirmation of *warA* deletion. Primer pair 1 was used to confirm deletion of *warA* ORF – deletion strains are negative, WT is positive. Primer pair 2 was used to confirm integration of *HygR* at the *warA* locus. (C) Southern blotting of *warA* and *cdcA/warA* deletion strains. gDNA of strains was digested with *Hin*dIII restriction enzyme. Membranes were hybridised with a probe consisting of Digoxigenin-UTP labelled *HygR*. Single bands confirm single integration of deletion cassette into the *A. niger* genome. Transformants Δ*warA_12* and ΔΔ*cdcA/warA_12* were used for experiments.

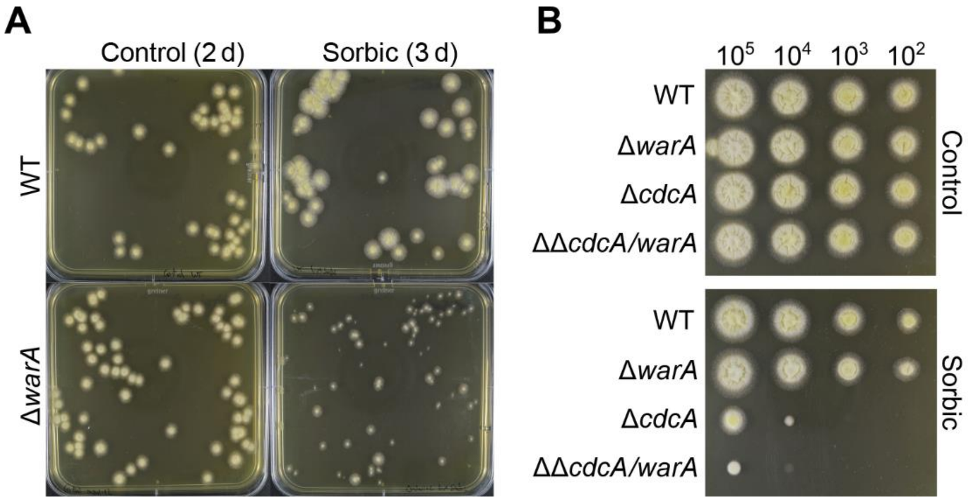


**FIG S3** (A) Growth of the Δ*warA* strain conidia on medium containing sorbic acid. Plates were inoculated with ~100 conidia and incubated for 2 d (control medium) or 3 d (medium containing 1 mM sorbic acid). (B) Radial growth of ΔΔ*cdcA/warA* strain on medium containing 1 mM sorbic acid. Plates were inoculated with a 10-fold dilution series of conidial suspensions; approximate numbers of conidia are indicated at the top. Images were captured after 2 d growth at 28°C, and are representative of 2-3 independent experiments.

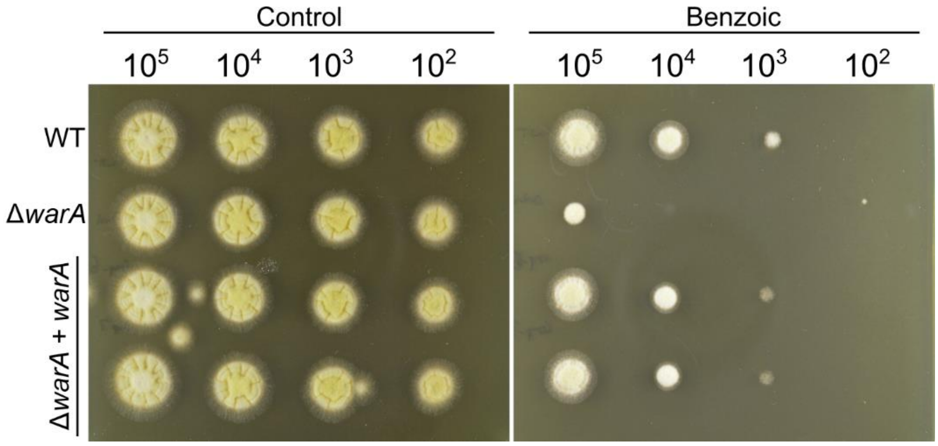


**FIG S4** Radial growth of complemented Δ*warA* strains on medium containing 2 mM benzoic acid. Plates were inoculated with a 10-fold dilution series of conidial suspensions. Approximate numbers of conidia are indicated above the pictures. Images were captured after 2 d growth at 28°C. Two independent complemented lines are shown.

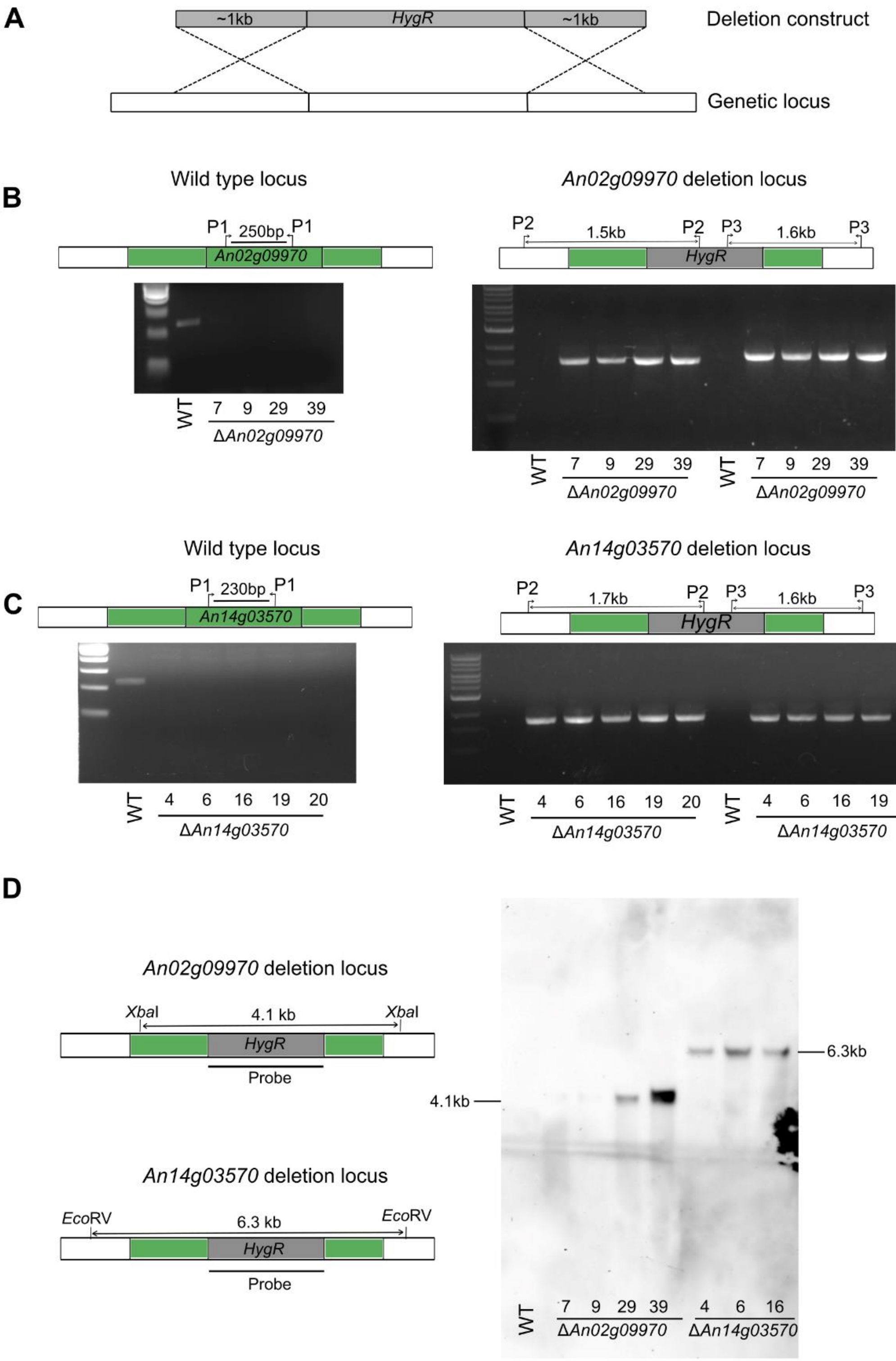


**FIG S5** PCR and Southern Blotting of *An02g09970* and *An14g03570* (*pdrA*) deletion strains. (A) Targeted gene deletion strategy. The *An02g09970* and *An14g03570* ORFs were replaced with a hygromycin resistance cassette. (B) PCR confirmation of *An02g09970* deletion. Primer pair 1 was used to confirm deletion of the *An02g09970* ORF – deletion strains are negative, WT is positive. Primer pairs 2 and 3 were used to confirm integration of *HygR* at the *An02g09970* locus. (C) PCR confirmation of *An14g03570* deletion. Primer pair 1 was used to confirm deletion of the *An14g03570* ORF – deletion strains are negative, WT is positive. Primer pairs 2 and 3 were used to confirm integration of *HygR* at the *An14g03570* locus. (D) Southern blotting of ΔA*n02g09970* and ΔA*n14g03570* deletion strains. gDNA of strains was digested with the restriction enzymes *Xba*I (for ΔA*n02g09970*) or *Eco*RV (for ΔA*n14g03570*). Membranes were hybridised with a probe consisting of Digoxigenin-UDP labelled *HygR*. Single bands confirm single integration of deletion cassette into the *A. niger* genome (multiple integrations apparent for Δ*An02g09970_39*). Transformants ΔA*n14g03570_4* and Δ*An02g09970_29* were used for experiments.

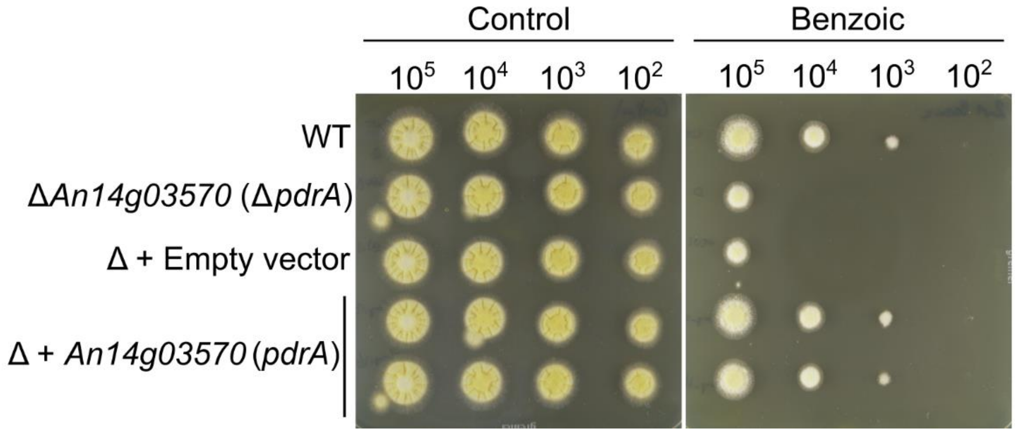


**FIG S6** Radial growth of complemented Δ*pdrA* strains on medium containing 2mM benzoic acid. Plates were inoculated with a 10-fold dilution series of conidial suspensions. Two independent complemented lines are shown. A Δ*pdrA* strain containing the empty pAN7.1*BAR* plasmid is also shown.

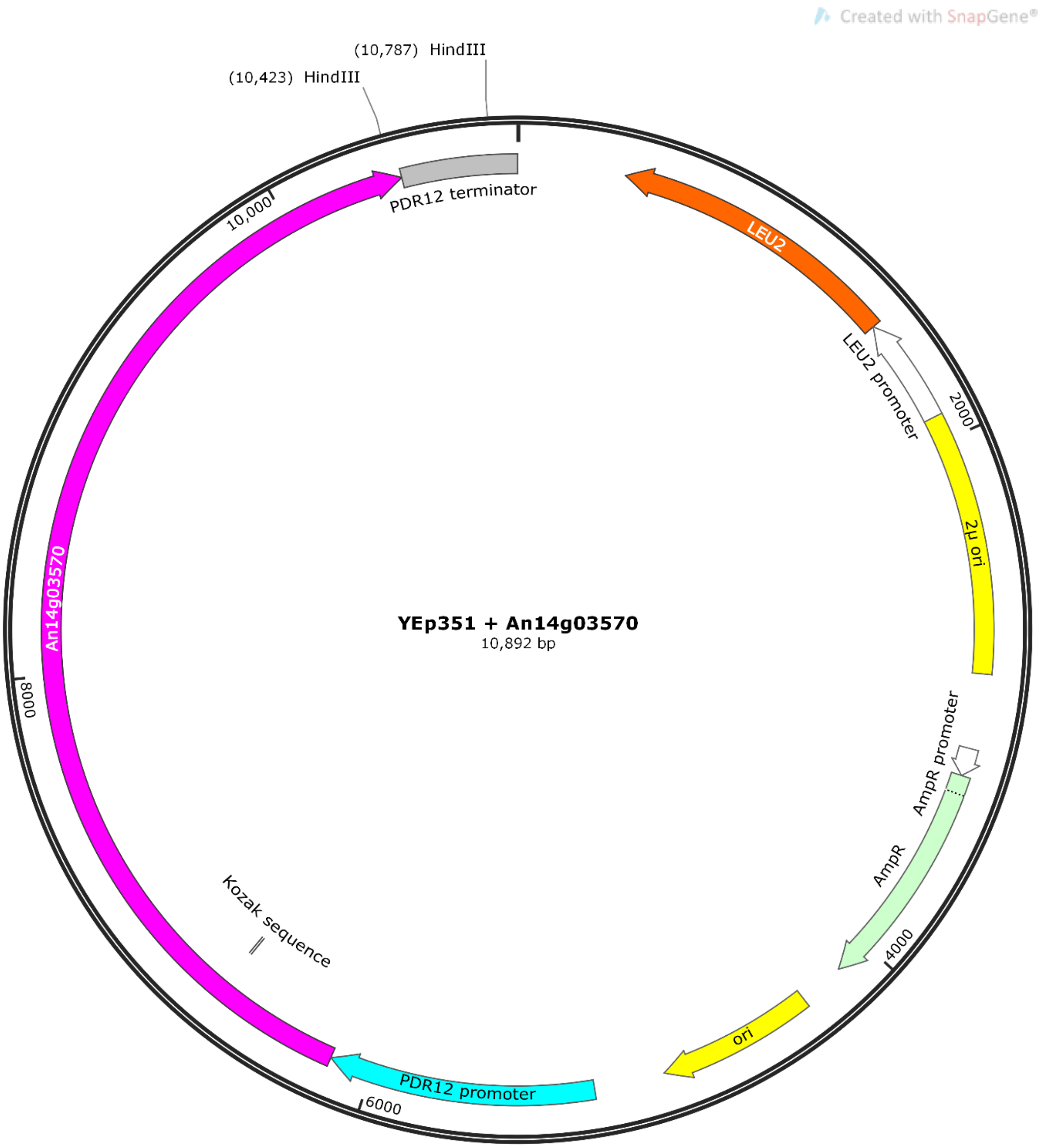


**FIG S7** Plasmid map of YEp351 containing *An14g03570* (*pdrA*)

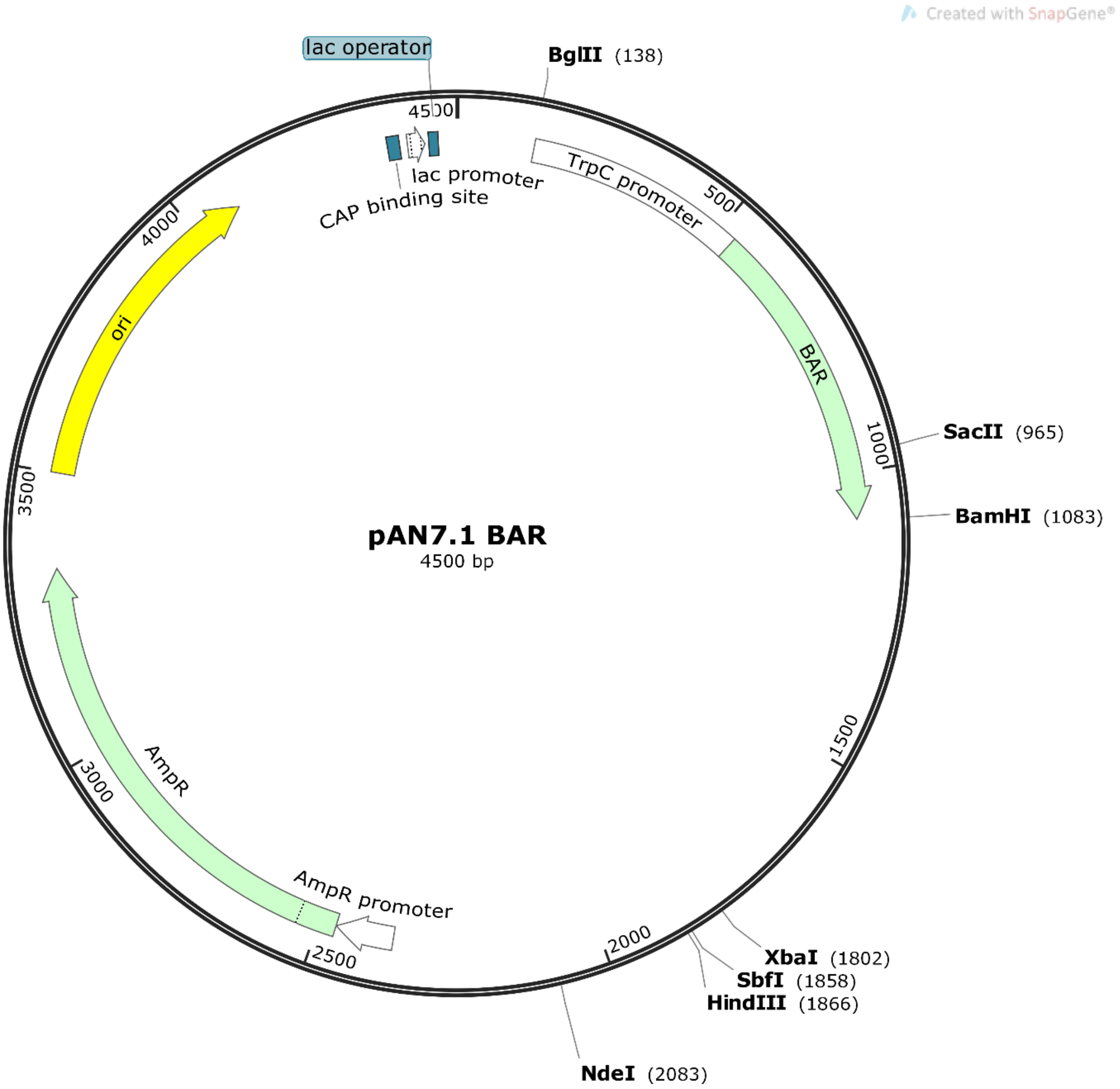


**FIG S8** Plasmid map of pAN7.1*BAR*.

**TABLE S1** RPKM and $Log_2FC$ values for *A. niger* genes (Excel file).

**TABLE S2** List of primers used in this study (Excel file).
